# Macrophages regulate gastrointestinal motility through complement component 1q

**DOI:** 10.1101/2022.01.27.478097

**Authors:** Mihir Pendse, Yun Li, Cristine N. Salinas, Gabriella Quinn, Nguyen Vo, Daniel C. Propheter, Chaitanya Dende, Alexander A. Crofts, Eugene Koo, Brian Hassell, Kelly A. Ruhn, Prithvi Raj, Yuuki Obata, Lora V. Hooper

**Affiliations:** Department of Immunology, The University of Texas Southwestern Medical Center, Dallas, TX 75390; The Howard Hughes Medical Institute, The University of Texas Southwestern Medical Center, Dallas, TX 75390

## Abstract

Peristaltic movement of the intestine propels food down the length of the gastrointestinal tract to promote nutrient absorption. Interactions between intestinal macrophages and the enteric nervous system regulate gastrointestinal motility, yet we have an incomplete understanding of the molecular mediators of this crosstalk. Here we identify complement component 1q (C1q) as a macrophage product that regulates gut motility. Macrophages were the predominant source of C1q in the mouse intestine and most extraintestinal tissues. Although C1q mediates complement-mediated killing of bacteria in the bloodstream, we found that C1q was not essential for immune defense of the intestine. Instead, C1q-expressing macrophages were localized to the intestinal submucosal plexus where they closely associated with enteric neurons and expressed surface markers characteristic of nerve-adjacent macrophages in other tissues. Mice with a macrophage-specific deletion of *C1qa* showed changes in enteric neuronal gene expression, increased peristaltic activity, and accelerated intestinal transit. Our findings identify C1q as a key regulator of gastrointestinal motility and provide enhanced insight into the crosstalk between macrophages and the enteric nervous system.

## INTRODUCTION

Peristalsis is the physical force that propels food through the intestine, promoting digestion and nutrient absorption. The gastrointestinal motility that underlies peristalsis is a complex process that requires coordination of the activity of smooth muscle cells by enteric neurons (Rao and Gershon, 2016). Recent studies have revealed that intestinal macrophages impact gastrointestinal motility by regulating the functions of enteric neurons and facilitating their interactions with smooth muscle cells (Muller et al., 2014; Matheis et al., 2020).

Macrophages carry out diverse functions in the intestine that vary according to their anatomic location. For example, macrophages that localize to the tissue located directly underneath the gut epithelium — known as the lamina propria — contribute to immune defense against pathogenic bacteria (Gabanyi et al., 2016). A distinct group of macrophages localizes to the tissues located beneath the lamina propria, between the circular and longitudinal muscle layers in the tissue region known as the muscularis externa. These muscularis macrophages express genes that are distinct from lamina propria macrophages (Gabanyi et al., 2016). They directly regulate the activity of smooth muscle cells (Luo et al., 2018) and secrete soluble factors such as bone morphogenetic protein 2 (BMP2) that interact with the enteric neurons that control smooth muscle activity (Muller et al., 2014). Muscularis macrophages thus play a key role in regulating gut motility. However, our understanding of the molecular mechanisms by which these macrophages regulate intestinal neuromuscular activity and gut motility remains limited.

C1q is a member of the defense collagen family that has distinct functions in immune defense and nervous system development and function (Bossi et al., 2014; Casals et al., 2019; Shah et al., 2015; Thielens et al., 2017). It is composed of six molecules each of C1qA, C1qB, and C1qC, forming a 410 kDa oligomer. C1q circulates in the bloodstream, where it participates in immune defense against infection by recognizing antibodies bound to invading bacteria. This binding interaction initiates the classical complement pathway, which entails the recruitment and proteolytic processing of other complement components that rupture the bacterial membrane and recruit phagocytic cells (Kishore and Reid, 2000; Noris and Remuzzi, 2013). C1q is also produced by microglia (brain-resident macrophage-like cells) in the brain where it promotes the pruning of neuronal synapses through an unclear mechanism (Hammond et al., 2020; Hong et al., 2016). Consequently, C1q deficiency results in heightened synaptic connectivity in the central nervous system that can lead to epilepsy (Chu et al., 2010).

C1q is also produced at barrier sites, such as the intestine, where encounters with commensal and pathogenic microbes are frequent. However, little is known about the physiological role of C1q in barrier tissues. Liver immune cells, including macrophages and dendritic cells, produce serum C1q; however the cellular source of C1q in barrier tissues including the intestine remains unclear (Petry et al., 2001). Here, we show that C1q is produced by macrophages that inhabit the submucosal plexus of the mouse intestine. These C1q-expressing macrophages are located close to enteric neurons that have a known role in controlling gut motility. Consistent with their nerve-adjacent localization, mice lacking macrophage C1q exhibit altered expression of enteric neuronal genes, increased peristaltic activity, and accelerated gastrointestinal motility. These findings identify C1q as a key mediator of a neuroimmune interaction that regulates gut motility.

## RESULTS

### C1q is expressed in intestinal macrophages

Soluble defense collagens are an ancient, evolutionarily conserved family of antimicrobial proteins with shared structural features including a *C*-terminal globular head and a collagen-like region (Casals et al., 2019). Little is known about the function of defense collagens at mucosal barrier sites, where microbial encounter is frequent. Our initial goal in this study was to identify soluble defense collagens that are expressed by the mouse intestine and to assess their role in host defense. We therefore measured the expression of 18 defense collagen genes in the mouse small intestine and colon by RNA sequencing (RNA-seq). The most abundant soluble defense collagen transcripts in the small intestine and colon were those encoding C1qA, C1qB, and C1qC (***Figure 1A***; ***Figure 1 – figure supplement 1***).

**Figure 1.**
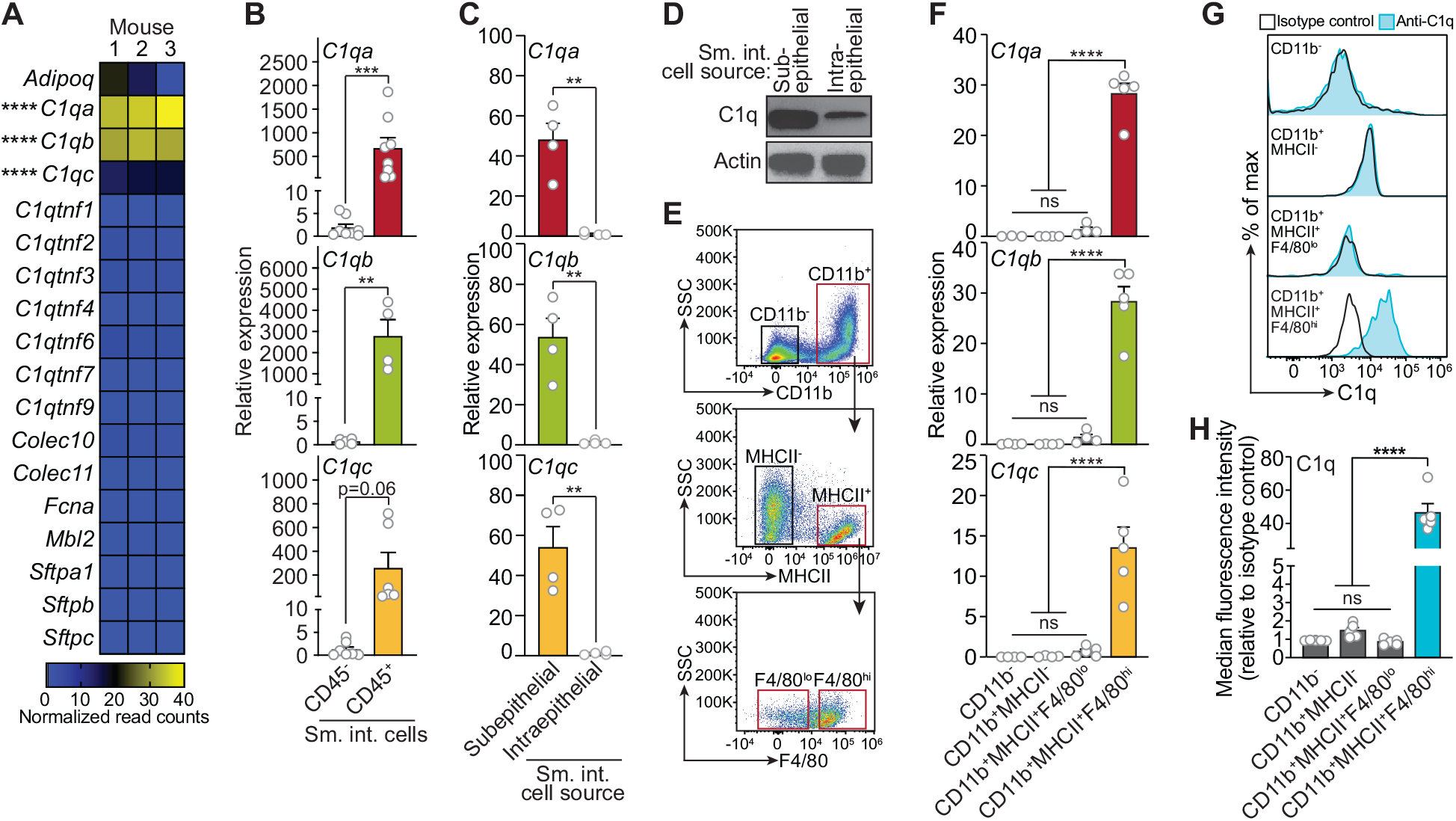
C1q is expressed by macrophages in the mouse small intestine. **(A)** RNA-seq analysis of soluble defense collagen expression in the small intestines (ileum) of C57BL/6 mice. Data were adapted from a previously published RNA-seq analysis (Gattu et al., 2019). Data are available in the Gene Expression Omnibus repository under accession number GSE122471. Each column represents one mouse. **(B)** qPCR measurement of *C1qa*, *C1qb*, and *C1qc* transcript abundance in CD45^+^ and CD45^-^ cells purified from mouse small intestines by flow cytometry. Each data point represents one mouse, and results are representative of two independent experiments. **(C)** qPCR measurement of *C1qa*, *C1qb*, and *C1qc* transcript abundance in subepithelial and intraepithelial cells recovered from mouse small intestines. Each data point represents one mouse, and results are representative of three independent experiments. **(D)** Representative immunoblot of subepithelial and intraepithelial cells recovered from mouse small intestines, with detection of C1q and actin (control). Each lane represents cells from one mouse and immunoblot is representative of three independent experiments. **(E)** Flow cytometry gating strategy for analysis of mouse small intestinal cell suspensions in panels F, G, and H. Cells were pre-gated as live CD45^+^ cells. SSC, side-scatter; MHCII, major histocompatibility complex II. **(F)** qPCR measurement of *C1qa*, *C1qb*, and *C1qc* transcript abundance in cells isolated by flow cytometry from mouse small intestines as indicated in (E). Each data point represents cells pooled from three mice, and results are representative of three independent experiments. **(G)** Flow cytometry analysis of intracellular C1q in small intestinal subepithelial cells identified as indicated in (E). **(H)** Quantitation of flow cytometry analysis in (G). Each data point represents one mouse, and results are representative of two independent experiments. Sm. int., mouse small intestine; Error bars represent SEM. **p<0.01; ***p<0.001; ****p<0.0001; ns, not significant by one way ANOVA (A,F) or two-tailed Student’s t-test (B,C,H). **Figure supplement 1.** C1q is expressed in the mouse colon.

Serum C1q is produced by liver dendritic cells, monocytes, and macrophages (El-Shamy et al., 2018). However, the cellular source(s) of C1q in peripheral tissues including the intestine is unknown. Quantitative PCR (qPCR) analysis of fluorescence-activated cell sorting (FACS)-sorted cell suspensions recovered from the small intestines of wild-type C57BL/6 mice revealed that *C1qa*, *C1qb*, and *C1qc* transcripts were most abundant in CD45^+^ cells, which include all immune cells, as compared to CD45^-^ cells, which encompass epithelial cells and other non-immune cells (***Figure 1B***). Further, C1q transcripts and protein were most abundant in CD45^+^ cells recovered from the subepithelial compartment, which includes both the lamina propria and muscularis, as compared to CD45^+^ cells recovered from the intraepithelial compartment of the small intestine (***Figure 1C and D***). Thus, C1q is expressed by immune cells located in the subepithelial compartment of the intestine and is largely absent from epithelial cells and intraepithelial immune cells.

To identify intestinal immune cells that expresse C1q, we further analyzed the subepithelial CD45^+^ cell population by flow cytometry. Expression of C1q transcripts and protein was highest in CD11b^+^MHCII^+^F4/80^hi^ macrophages and was mostly absent from non-macrophage immune cells (***Figure 1E-H***). Thus, C1q is expressed by macrophages in the mouse small intestine.

### Macrophages are the primary source of C1q in the mouse gastrointestinal tract

We next assessed whether macrophages are the primary source of C1q in the intestine by analyzing two mouse models. First, we depleted macrophages by injecting neutralizing antibodies directed against the receptor for colony stimulating factor 1 (CSF1R)(***Figure 2A***), which is required for the development of a subset of lamina propria macrophages (Bogunovic et al., 2009) and all muscularis macrophages (Muller et al., 2014). Antibody injection led to a >2-fold reduction in the number of macrophages recovered from the small intestine (***Figure 2B***), and a corresponding >2-fold reduction in small intestinal C1q gene expression (***Figure 2C***), suggesting that macrophages are the primary source of intestinal C1q.

**Figure 2.**
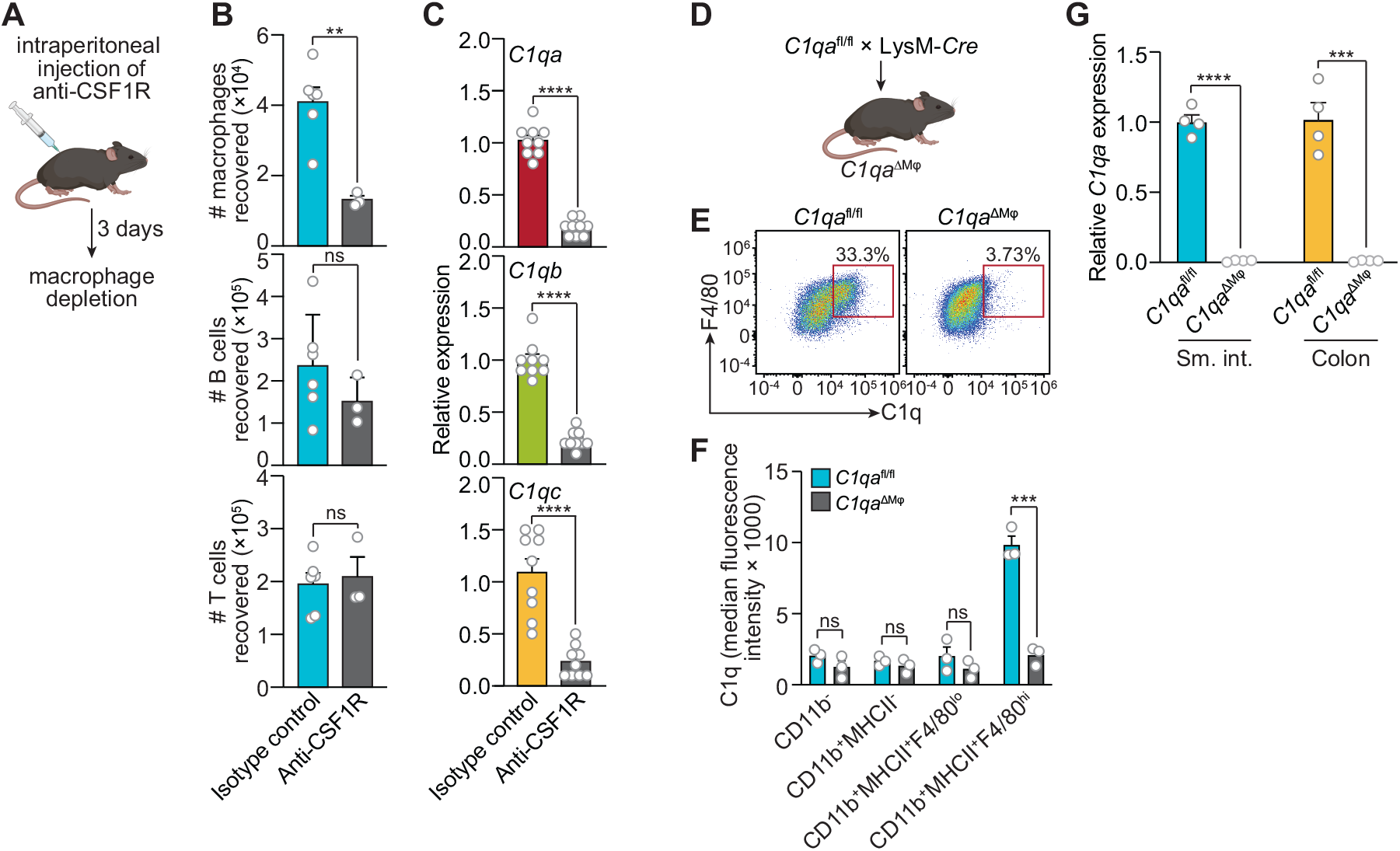
Macrophages are the primary source of C1q in the mouse gastrointestinal tract. **(A)** Graphic showing selective depletion of macrophages in C57BL/6 mice by intraperitoneal injection of anti-CSF1R antibody. Control mice were injected with isotype-matched non-specific antibody. Mice were analyzed 72 hours after antibody injection. Panel was generated at Biorender.com. **(B)** Representative flow cytometry analysis of mouse small intestines after intraperitoneal injection of anti-CSF1R or isotype control antibody. All cells were gated as live CD45^+^. Macrophages were MHCII^+^ F4/80^hi^; B cells were CD19^+^; T cells were CD3^+^. Total small intestinal cell yields were 1.5 × 10^6^ ± 4.9 × 10^5^ cells. **(C)** qPCR measurement of *C1qa*, *C1qb*, and *C1qc* transcript abundance in mouse small intestines after intraperitoneal injection of anti-CSF1R or rat IgG2a (isotype control). Each data point represents one mouse and results are pooled from two independent experiments. **(D)** *C1qa*^fl/fl^ mice were crossed with mice harboring a LysM-Cre transgene to generate mice having a macrophage-selective deletion of *C1qa* (*C1qa*^ΔMφ^ mice). Panel was generated at Biorender.com. **(E)** Representative flow cytometry analysis of intracellular C1q expression in small intestinal macrophages from *C1qa*^fl/fl^ and *C1qa*^ΔMφ^ mice. Mice were littermates from heterozygous crosses that remained co-housed. Cells were gated on live CD45^+^CD11b^+^MHCII^+^. **(F)** Quantitation of the flow cytometry analysis in (E). Each data point represents one mouse. Results are representative of two independent experiments. **(G)** qPCR measurement of *C1qa* transcript abundance in the small intestines (sm. int.) and colons of *C1qa*^fl/fl^ and *C1qa*^ΔMφ^ littermates. Each data point represents one mouse. Error bars represent SEM. **p<0.01; ***p<0.001; ****p<0.0001; ns, not significant by two-tailed Student’s t-test. **Figure supplement 1.** C1q expression is lost systemically but preserved in the central nervous system of *C1qa*^ΔMφ^ mice.

Second, we constructed a genetic model of C1q deficiency by crossing *C1qa*^fl/fl^ mice (Fonseca et al., 2017) to mice carrying the *Lyz2-Cre* transgene (LysM-Cre mice), which is selectively expressed in myeloid cells including macrophages (***Figure 2D***). These mice, hereafter designated as *C1qa*^ΔMϕ^ mice, lacked C1q expression in intestinal macrophages (***Figure 2E and F***). Importantly, *C1qa*^ΔMϕ^ mice had markedly lower C1q expression in both the small intestine and colon (***Figure 2G***), indicating that macrophages are the main source of C1q in the intestine. Unexpectedly, the *C1qa*^ΔMϕ^ mice also lost *C1q* gene expression in the lung, skin, kidney, and liver (but not the brain), and C1q protein was undetectable in the serum (***Figure 2 – figure supplement 1***). These findings indicate that macrophages are the primary source of C1q in the intestine and suggest that LysM^+^ macrophages or macrophage-like cells are also the main source of C1q in most extraintestinal tissues and the bloodstream.

### *C1qa*^ΔMφ^ mice do not show altered microbiota composition, barrier function, or resistance to enteric infection

The classical complement pathway is a well-studied host defense system that protects against systemic pathogenic infection (Warren et al., 2002; Noris and Remuzzi, 2013). Circulating C1q activates the complement pathway by binding to antibody-antigen complexes or to bacterial cell surface molecules, and thus protects against systemic infection. We therefore assessed whether C1q promotes immune defense of the intestine.

We first determined whether C1q exhibits characteristics of known intestinal antimicrobial proteins, including induction by the intestinal microbiota and secretion into the gut lumen. *C1qa* was expressed at similar levels in the small intestines of germ-free and conventionally-raised mice (***Figure 3A***), suggesting that *C1q* expression is not induced by the gut microbiota. This contrasted with *Reg3g*, encoding the antimicrobial protein REG3G (Cash et al., 2006), which was expressed at a >2-fold higher level in conventional as compared to germ-free mice (***Figure 3A***). Additionally, in contrast to REG3G, C1q was not detected in the gut lumen of either conventional or germ-free mice (***Figure 3B***). *C1qa* expression was also not markedly altered by oral infection with the intestinal pathogenic bacterial species *Salmonella* Typhimurium (***Figure 3C***). These data indicate that C1q is not induced by the gut microbiota or by the bacterial pathogen *S.* Typhimurium, in contrast to other antibacterial proteins.

**Figure 3.**
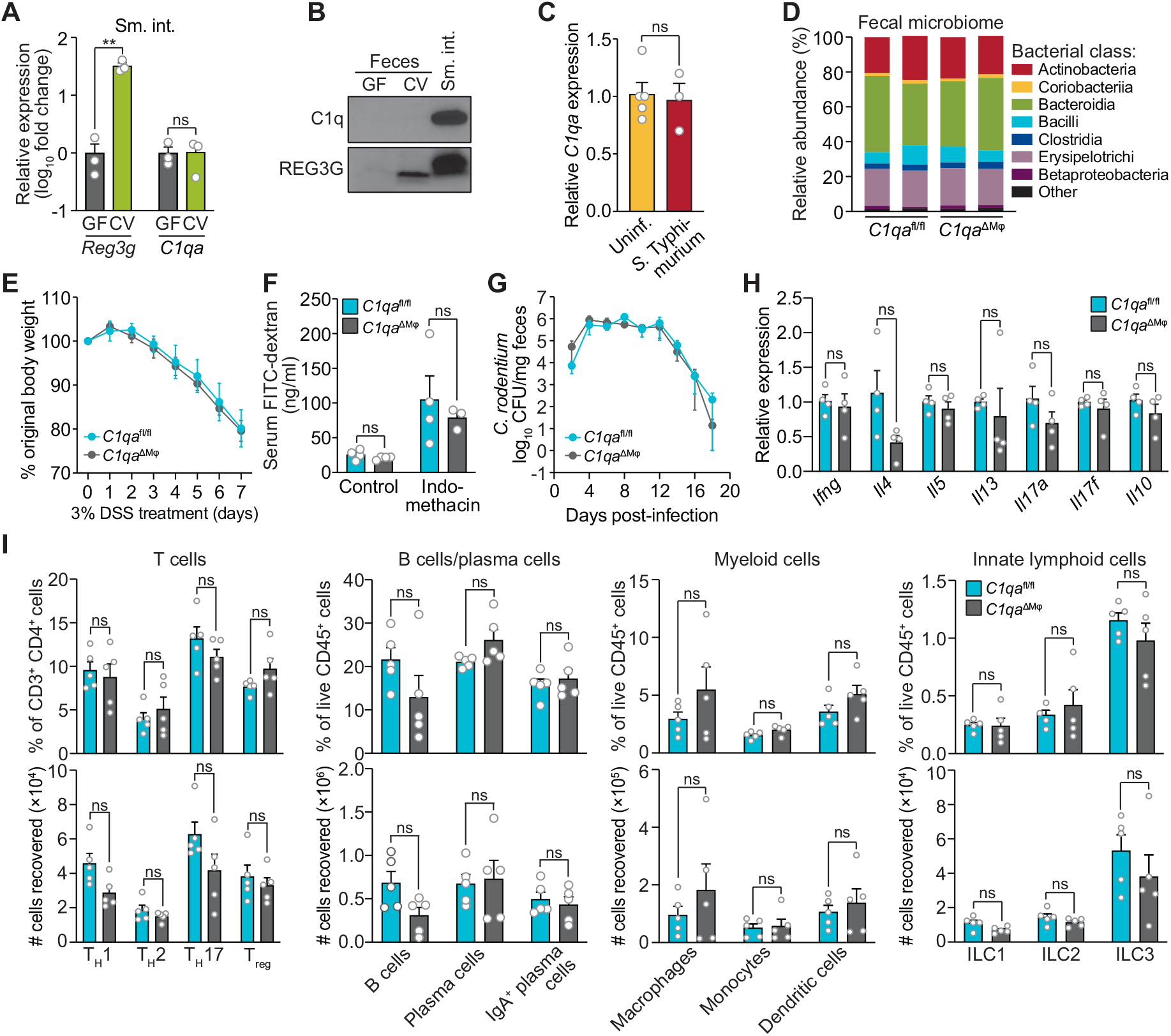
*C1qa*^ΔMφ^ mice do not show altered microbiota composition, barrier function, or resistance to enteric infection. **(A)** Small intestinal *C1qa* expression is not induced by the intestinal microbiota. qPCR measurement of *Reg3g* and *C1qa* transcript abundances in the small intestines of germ free (GF) and conventional (CV) C57BL/6 mice. Each data point represents one mouse and results are representative of two independent experiments. **(B)** C1q is not detected in the mouse intestinal lumen or feces. Representative immunoblot of an ammonium sulfate precipitation of intestinal lumenal contents and feces from germ-free and conventional mice with detection of C1q. C1q in small intestinal tissue (sm. int.) is shown for comparison at right. REG3G was detected as a control, as it is secreted into the intestinal lumen of conventional mice (Cash et al., 2006). Each lane represents multiple mice pooled (n = 5 and 9 for germ-free and conventional, respectively) and the immunoblot is representative of three independent experiments. **(C)** C1q gene expression is not altered by enteric infection with *S.* Typhimurium. qPCR measurement of *C1qa* transcript abundance in small intestinal tissue after oral inoculation of mice with 10^9^ colony forming units of *S.* Typhimurium strain SL1344. Each data point represents one mouse, and results are representative of two independent experiments. **(D)** Intestinal microbiota composition is not altered in *C1qa*^ΔMφ^ mice. Phylogenetic analysis of 16*S* rRNA gene sequences from fecal pellets collected from *C1qa*^fl/fl^ and *C1qa*^ΔMφ^ littermates. Operational taxonomic units with an average of 100 reads and populations greater than or equal to 1% were included in the graphical analysis. Each bar represents one mouse. Data are available from the Sequence Read Archive under BioProject ID PRJNA793870. **(E)** *C1qa*^ΔMφ^ mice do not show altered susceptibility to dextran sulfate sodium (DSS)-induced colitis. Mice were provided with 3% DSS in drinking water and body weights were monitored for 7 days. n = 4 and 6 for *C1qa*^fl/fl^ and *C1qa*^ΔMφ^ littermates, respectively. Error bars represent SEM. Differences at each time point were not significant by two-tailed Student’s t-test. **(F)** *C1qa*^ΔMφ^ mice do not show altered intestinal permeability. To measure intestinal permeability, *C1qa*^fl/fl^ and *C1qa*^ΔMφ^ littermates were gavaged with FITC-dextran, and serum FITC-dextran levels were determined by fluorescence microplate assay against a FITC-dextran standard curve. Indomethacin induces intestinal damage in mice and was used as a positive control. **(G)** Time course of fecal *C. rodentium* burden following oral gavage of *C1qa*^fl/fl^ and *C1qa*^ΔMφ^ mice with 5 × 10^8^ colony forming units (CFU) of *C. rodentium*. n = 5 and 5 for *C1qa*^fl/fl^ and ^C1qaΔMφ^ littermates, respectively. Error bars represent SEM. Differences at each time point were not significant by two-tailed Student’s t-test. **(H)** qPCR measurement of cytokine transcript abundances in the small intestines of *C1qa*^fl/fl^ and *C1qa*^ΔMφ^ littermates. Error bars represent SEM. Statistics were performed with two-tailed Student’s t-test. **(I)** Flow cytometry analysis of small intestinal immune cell subsets from *C1qa*^fl/fl^ and *C1qa*^ΔMφ^ littermates. Gating strategies are shown in Figure 3 - figure supplement 1 through 4. ILC, innate lymphoid cell. Total small intestinal cell yields were 8.8 × 10^6^ ± 2.9 × 10^6^ cells. Error bars represent SEM. **p<0.01; ns, not significant by two-tailed Student’s t-test. **Figure supplement 1.** Flow cytometry gating strategy for comparison of T cell populations in *C1qa*^fl/fl^ and *C1qa*^ΔMφ^ mice. **Figure supplement 2.** Flow cytometry gating strategy for comparison of B cell and plasma cell populations in *C1qa*^fl/fl^ and *C1qa*^ΔMφ^ mice. **Figure supplement 3.** Flow cytometry gating strategy for comparison of myeloid cell populations in *C1qa*^fl/fl^ and *C1qa*^ΔMφ^ mice. **Figure supplement 4.** Flow cytometry gating strategy for comparison of innate lymphoid cell populations in *C1qa*^fl/fl^ and *C1qa*^ΔMφ^ mice.

We next assessed whether C1q regulates the composition of the gut microbiota. 16*S* rRNA gene sequencing analysis of the fecal microbiotas of *C1qa*^fl/fl^ and *C1qa*^ΔMϕ^ mice showed that the microbiota composition was not appreciably altered in the absence of macrophage C1q (***Figure 3D***). We also challenged *C1qa*^fl/fl^ and *C1qa*^ΔMϕ^ mice with dextran sulfate sodium (DSS), which damages the colonic epithelium and exposes underlying tissues to the commensal microbiota. However, the sensitivity of the *C1qa*^ΔMϕ^ mice to DSS was similar to that of their *C1qa*^fl/fl^ littermates as assessed by change in body weight (***Figure 3E***). There was also no change in intestinal paracellular permeability in *C1qa*^ΔMϕ^ mice as measured by oral administration of FITC-dextran (***Figure 3F***). These results suggest that macrophage C1q does not substantially impact gut microbiota composition or intestinal epithelial barrier function.

To determine whether C1q protects against enteric infection we conducted oral infection experiments with the enteric pathogen *Citrobacter rodentium*. We chose *C. rodentium* as our model organism for two reasons. First, *C. rodentium* is a non-disseminating pathogen, allowing us to test specifically for C1q’s role in intestinal infection. Second, *C. rodentium* clearance depends on immunoglobulins and complement component C3 (Belzer et al., 2011). Because C1q is bactericidal in concert with C3 and immunoglobulins, we predicted that *C1qa*^ΔMϕ^ mice would be more susceptible to *C. rodentium* infection. However, *C1qa*^ΔMϕ^ mice cleared *C. rodentium* similarly to their *C1qa*^fl/fl^ littermates, indicating that C1q is dispensable for defense against *C. rodentium* infection (***Figure 3G***).

We also did not observe marked alterations in immunity in the absence of C1q. Comparison of cytokine transcript abundance in the small intestines of *C1qa*^fl/fl^ and *C1qa*^ΔMϕ^ littermates revealed no statistically significant differences in cytokine expression (***Figure 3H***). There were also no statistically significant differences in the percentages or absolute numbers of various T cell subsets, including T_helper_1 (T_H_1), T_H_2, T_H_17, and regulatory T (T_reg_) cells between *C1qa*^fl/fl^ and *C1qa*^ΔMϕ^ mice (***Figure 3I***; ***Figure 3 – figure supplement 1***). Although total B cell numbers trended lower in *C1qa*^ΔMϕ^ mice, the difference was not statistically significant (***Figure 3I***; ***Figure 3 – figure supplement 2***). There were also no statistically significant differences in the percentages or absolute numbers of total plasma cells (***Figure 3I***; ***Figure 3 – figure supplement 2***), IgA^+^ plasma cells (***Figure 3I***; ***Figure 3 – figure supplement 2***), myeloid cells (***Figure 3I***; ***Figure 3 – figure supplement 3***), or innate lymphoid cells (***Figure 3I***; ***Figure 3 – figure supplement 4***) when comparing *C1qa*^fl/fl^ and *C1qa*^ΔMϕ^ mice. These results suggest that the absence of macrophage C1q has little impact on intestinal immunity. Altogether, our findings suggest that C1q does not participate substantially in intestinal immune defense and thus might have an intestinal function that is independent of its canonical role in activating the classical complement pathway.

### C1q-expressing gut macrophages are located near enteric neurons

Intestinal macrophages perform distinct functions depending on their anatomical location. Macrophages in the lamina propria protect against invasion by pathogenic microbes and promote tissue repair (Grainger et al., 2017). In contrast, macrophages that reside in deeper intestinal tissues, including the submucosal plexus and muscularis (***Figure 4A***), regulate enteric smooth muscle cells and neurons that drive gastrointestinal motility (De Schepper, Stakenborg, et al., 2018; De Schepper, Verheijden, et al., 2018).

**Figure 4.**
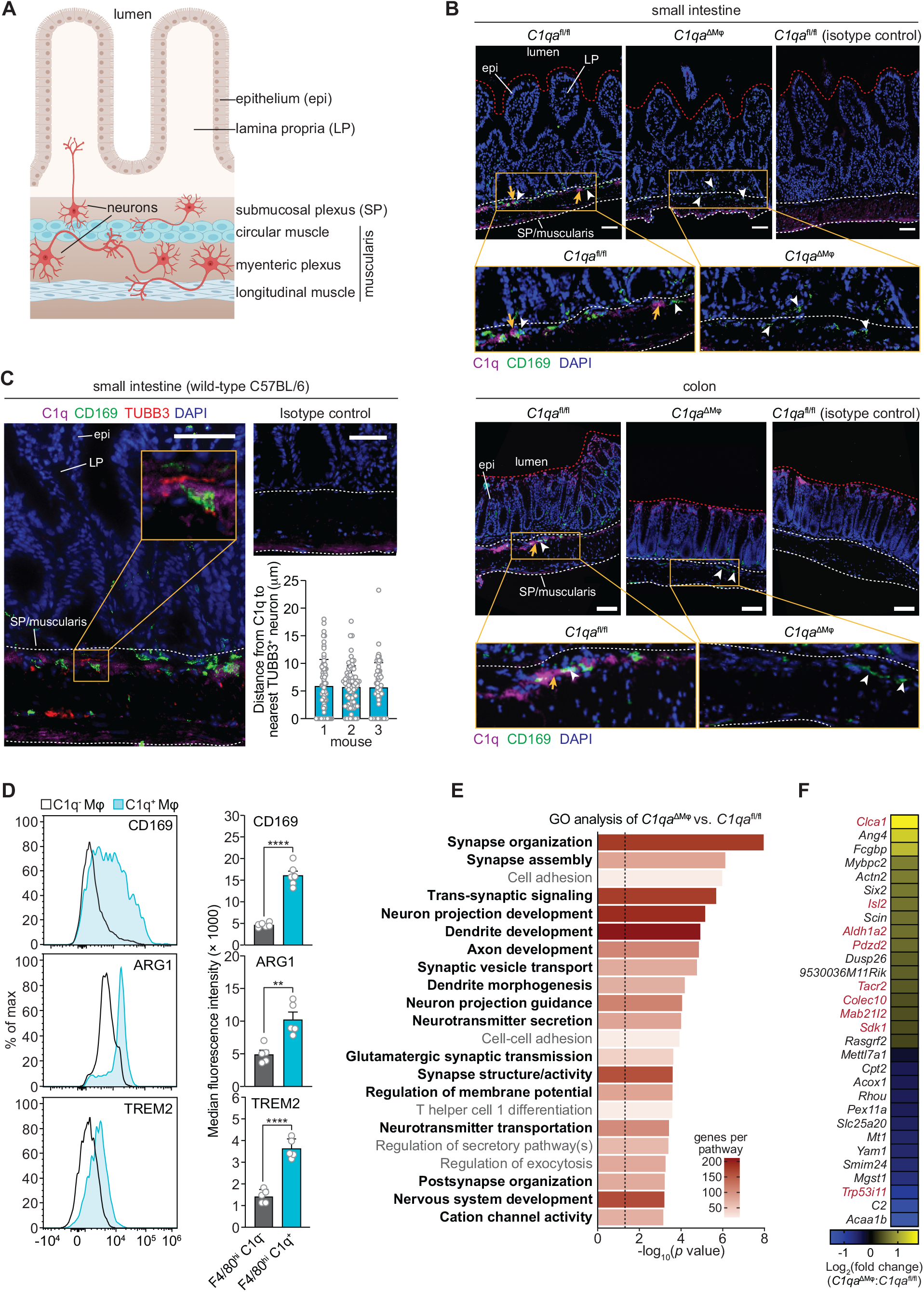
C1q-expressing macrophages are closely associated with neurons in the mouse intestine. **(A)** Graphic depicting the muscularis of the mouse small intestine. The lumen, epithelium (epi), lamina propria (LP), submucosal plexus (SP), and distinct anatomical regions of the muscularis are indicated. Created at Biorender.com. **(B)** Immunofluorescence detection of C1q (violet) and macrophages marked with CD169 (green) in the small intestine and colon of *C1qa*^fl/fl^ and *C1qa*^ΔMφ^ littermates. Nuclei were detected with 4’,6-diamidino-2-phenylindole (DAPI; blue). Detection with isotype control antibodies on *C1qa*^fl/fl^ small intestines is shown at right. Anti-rat IgG AlexaFluor 488 and streptavidin-Cy5 were used as secondary stains for CD169 and C1q respectively. The intestinal surface is denoted with a red dotted line and the gut lumen, epithelium, and lamina propria are indicated. The approximate region encompassing the submucosal plexus and the muscularis is denoted with two white dotted lines. Examples of C1q^+^ areas are indicated with yellow arrows and examples of CD169^+^ macrophages are indicated with white arrowheads. Note that the violet staining near the bottom of the muscularis is non-specific, as indicated by its presence in the isotype control image. Images are representative of three independent experiments. Scale bars = 50 μm. **(C)** Immunofluorescence detection of C1q (violet), macrophages marked with CD169 (green) and neurons marked with TUBB3 (red) in the small intestines of wild-type C57BL/6 mice. Nuclei are detected with DAPI (blue). The epithelium, lamina propria are indicated. The approximate region encompassing the submucosal plexus and the muscularis is denoted with two white dotted lines. The expanded image area delineated by a yellow square shows an example of the close association between C1q and TUBB3^+^ neurons. Images are representative of images captured from three mice. Anti-rat IgG AlexaFluor 488, anti-rabbit IgG AlexaFluor 594, and streptavidin-Cy5 were used as secondary stains for CD169, TUBB3, and C1q respectively, and an isotype control image is shown at upper right. Scale bars = 50 μm. Lower right panel: Distances between C1q^+^ areas and the nearest TUBB3^+^ neurons were measured in images captured from three mice. Each data point represents one measurement and measurements were made on 10 to 15 microscopic fields for each mouse (71, 70, and 57 total measurements, respectively). **(D)** Flow cytometry analysis of CD169, Arginase 1, and TREM2 on C1q^-^ and C1q^+^ macrophages recovered from the small intestines of wild-type C57BL/6 mice. Median fluorescence intensities from multiple mice are quantified in the panels at right. Each data point represents one mouse (n=5-6 mice), and results are representative of two independent experiments. Error bars represent SEM. **p<0.01; ****p<0.0001 by two-tailed Student’s t-test. **(E)** Colonic longitudinal muscle myenteric plexus was acquired from *C1qa*^ΔMφ^ and *C1qa*^fl/fl^ littermates. RNA-seq was performed, and annotated gene ontology (GO) biological processes were assigned to genes that were differentially expressed in *C1qa*^ΔMφ^ mice when compared to their *C1qa*^fl/fl^ littermates. GO biological processes associated with neurons are in bold type. The dotted line indicates the cutoff for statistical significance. Five mice per group were analyzed as pooled biological replicates. Data are available from the Sequence Read Archive under BioProject ID PRJNA793870. **(F)** The colonic longitudinal muscle myenteric plexus of *C1qa*^ΔMφ^ mice has a transcriptional profile like that of mice with a gastrointestinal motility disorder. RNA-seq was performed on the colonic longitudinal muscle-myenteric plexus from five *C1qa*^fl/fl^ and five *C1qa*^ΔMφ^ littermates. Genes that were differentially expressed are represented in a heatmap that depicts log_2_(fold change). Genes that also showed altered expression in the TashT mouse line, which is a model of human Hirschsprung’s disease (Bergeron et al., 2015), are indicated in red. Statistical significance of overlap between differentially expressed genes in *C1qa*^ΔMφ^ and TashT mice was determined by Fisher’s exact test (p = 0.0032). **Figure supplement 1.** Similar numbers of CD169^+^ macrophages are recovered from the small intestines of *C1qa*^ΔMφ^ and *C1qa*^fl/fl^ littermates.

Immunofluorescence analysis revealed that C1q was located primarily beneath the lamina propria, in a region consistent with the location of the submucosal plexus of the small intestine and colon (***Figure 4A and B***). C1q was detected in a diffuse pattern close to macrophages in *C1qa*^fl/fl^ mice and was absent in *C1qa*^ΔMϕ^ mice (***Figure 4B***) despite the presence of similar overall numbers of CD169^+^ macrophages (***Figure 4 – figure supplement 1***). Although we did not observe direct colocalization of C1q with CD169, a macrophage marker that selectively marks nerve-adjacent macrophages (Ural et al., 2020), the pattern was consistent with prior observations of C1q deposition in the extracellular matrix in the arterial wall (Cao et al., 2003)(***Figure 4B***).

We next assessed the location of C1q relative to enteric neurons in the mouse small intestine. Detection of the neuronal marker βIII tubulin (TUBB3) revealed the presence of two distinct layers of enteric neurons: one just beneath the lamina propria that was consistent with the location of the submucosal plexus, and one in the muscularis that was consistent with the location of the myenteric plexus (***Figure 4C***). Immunofluorescence detection of C1q alongside TUBB3 revealed that C1q was located within an average of ∼5 μm of the nearest TUBB3^+^ enteric neurons in the submucosal plexus and was largely absent from the myenteric plexus (***Figure 4C***). Thus, C1q-producing gut macrophages are located close to enteric neurons in the submucosal plexus.

C1q-producing intestinal macrophages exhibited several characteristics of nerve-adjacent macrophages in other tissues. First, microglia in the central nervous system and nerve-adjacent macrophages in peripheral tissues, such as the lung, are enriched for C1q gene expression (Fonseca et al., 2017; Ural et al., 2020). This is consistent with our observations of C1q expression in nerve-adjacent intestinal macrophages located in the submucosal plexus (***Figure 4C***). Second, nerve-adjacent macrophages depend on a CSF1-CSF1R signaling axis for their maintenance and are therefore sensitive to anti-CSF1R antibody-mediated depletion (Muller et al., 2014). Thus, our finding that anti-CSF1R administration depleted C1q-expressing intestinal macrophages is consistent with their location near enteric neurons (***Figure 2A-C***). Third, C1q-expressing intestinal macrophages showed elevated expression of Arginase 1, CD169, and TREM2 (triggering receptor expressed on myeloid cells 2)(***Figure 4D***), which are enriched on macrophages with known neuromodulatory functions (Colonna, 2003; Paloneva et al., 2002; Ural et al., 2020). Thus, C1q-expressing intestinal macrophages are located near enteric neurons and exhibit phenotypes characteristic of nerve-adjacent macrophages in other tissues.

### *C1qa*^ΔMϕ^ mice have altered gastrointestinal motility

Macrophages engage in crosstalk with the enteric nervous system and regulate functions, including gastrointestinal motility, that depend on the enteric nervous system (Muller et al., 2014). This crosstalk involves the exchange of specific proteins such as bone morphogenetic protein 2 (BMP2) (Muller et al., 2014). Given that C1q^+^ macrophages phenotypically resemble peripheral nerve-adjacent macrophages and reside near enteric neurons, we postulated that macrophage-derived C1q might also regulate enteric nervous system function.

As an initial test of this idea, we performed RNA-seq on the colonic longitudinal muscle-myenteric plexus from *C1qa*^ΔMϕ^ and *C1qa*^fl/fl^ littermates and then conducted unbiased Gene Set Enrichment Analysis. Of the 22 biological pathways that were enriched in *C1qa*^ΔMϕ^ mice, 17 were related to neuronal development or function, including synapse organization, dendrite development, and neurotransmitter secretion (***Figure 4E***). 30 genes were differentially expressed when comparing the muscle-myenteric plexus of *C1qa*^ΔMϕ^ and *C1qa*^fl/fl^ mice (***Figure 4F***). These included genes with known roles in regulating neuronal activity (*Dusp26*), synaptic transmission (*Rasgrf2*), and neuropeptide signaling (*Tacr2*) (Mao et al., 2017; Schwechter et al., 2013; Yang et al., 2017). We also compared the list of genes differentially expressed in the *C1qa*^ΔMϕ^ mice to those differentially expressed in the *TashT* mouse line. This line contains an insertional mutation that leads to dysregulated gut motility comparable to Hirschsprung’s disease, a human genetic disorder resulting in incomplete development of the enteric nervous system (Bergeron et al., 2015). There was a statistically significant overlap in the transcriptional changes in the colonic longitudinal muscle-myenteric plexus of *C1qa*^ΔMϕ^ mice and *TashT* mice (***Figure 4F***). Together, these results suggest that macrophage C1q impacts enteric nervous system gene expression and function.

Efficient coordination of gastrointestinal motility is necessary for proper digestion, nutrient absorption, and excretion. Given that muscularis macrophages regulate enteric nervous system functions that govern gastrointestinal motility (Muller et al., 2014), we assessed whether macrophage C1q impacts gut motility. We first tested this idea by measuring gut transit time using the nonabsorbable dye carmine red. *C1qa*^ΔMϕ^ and *C1qa*^fl/fl^ littermates were gavaged with the dye and the time to first appearance of the dye in the feces was recorded. Transit times were decreased in *C1qa*^ΔMϕ^ mice relative to their *C1qa*^fl/fl^ littermates, indicating accelerated gut motility (***Figure 5A***). This was not due to a change in the lengths of either the small intestine or the colon, which were unaltered in the *C1qa*^ΔMϕ^ mice (***Figure 5B***). By contrast, gut transit time was unaltered in *C3^-/-^* mice, suggesting that macrophage C1q impacts gut motility independent of its canonical function in the classical complement pathway (***Figure 5A***). Accelerated transit was also observed in the small intestines of *C1qa*^ΔMϕ^ mice as assessed by rhodamine dye transit assay (***Figure 5C***). We then conducted *ex vivo* colonic peristaltic recordings and observed that the colons of *C1qa*^ΔMϕ^ mice exhibited greater neuromuscular activity as compared to *C1qa*^fl/fl^ littermates (***Figure 5D***; ***Figure 5 – figure supplement 1 and 2***). This indicated that the colons of *C1qa*^ΔMϕ^ mice maintained increased peristaltic activity compared to their *C1qa*^fl/fl^ littermates even in the absence of gut-extrinsic signals. Finally, we directly assessed colonic motility by bead expulsion assay and found that *C1qa*^ΔMϕ^ mice had accelerated colonic transit *in vivo* (***Figure 5E***). Thus, the absence of macrophage C1q increases peristaltic activity and accelerates gut transit, indicating that macrophage C1q regulates gastrointestinal motility.

**Figure 5.**
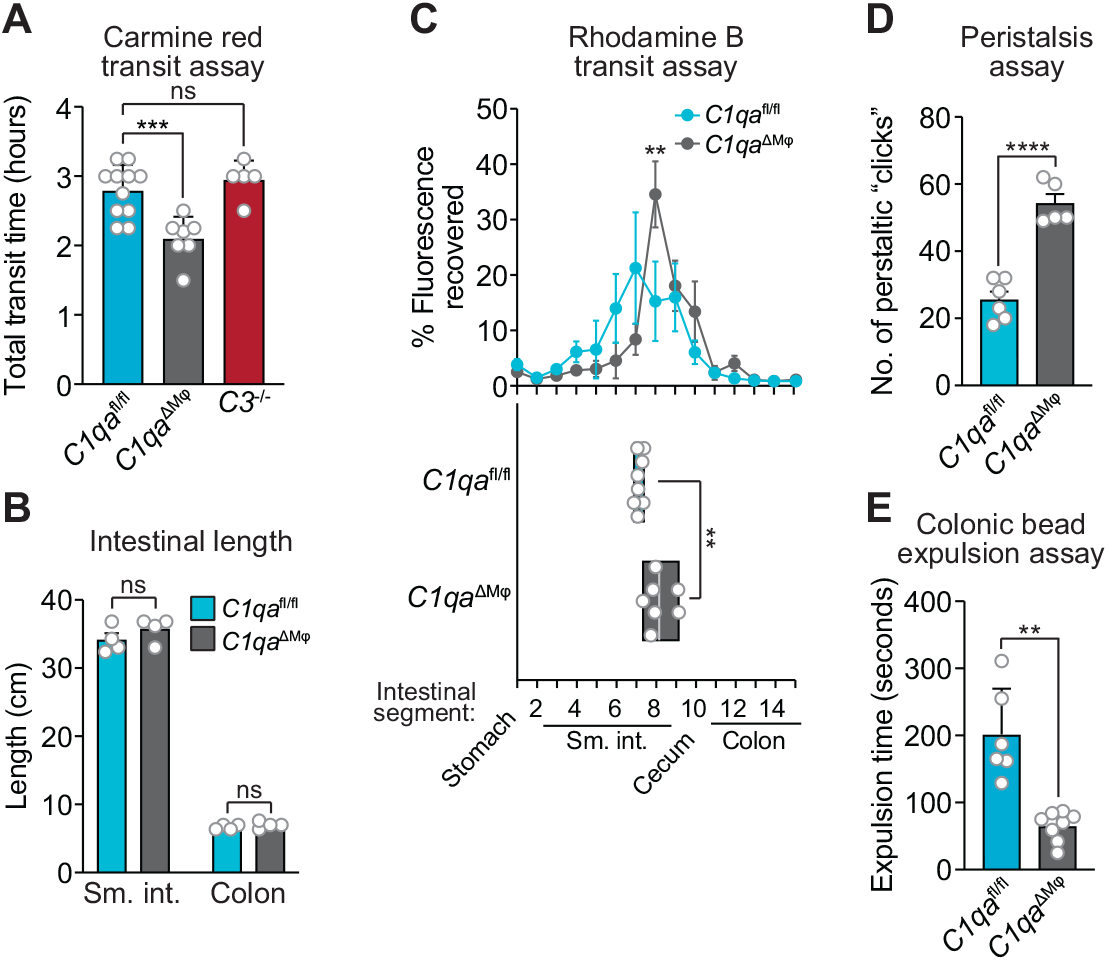
*C1qa*^ΔMφ^ mice have altered gastrointestinal motility. **(A)** Measurement of total intestinal transit time in *C1qa*^fl/fl^ and *C1qa*^ΔMφ^ littermates and *C3*^-/-^ mice. Mice were gavaged with 100 μl of carmine red (5% (w/v in 1.5% methylcellulose). Fecal pellets were collected every 15 minutes and transit time was recorded when the dye was first observed in the feces. Each data point represents one mouse and the results are pooled from five independent experiments. **(B)** Intestinal tract length is not altered in *C1qa*^ΔMφ^ mice. Small intestines and colons from *C1qa*^fl/fl^ and *C1qa*^ΔMφ^ littermates were excised and measured. Each data point represents one mouse. **(C)** Transit of rhodamine B-dextran through the intestines of *C1qa*^fl/fl^ and *C1qa*^ΔMφ^ littermates. Mice were sacrificed 90 minutes after gavage with rhodamine B-dextran. The intestines were divided into 16 segments, the rhodamine B fluorescence was measured in each segment (top panel), and the geometric center of the fluorescence was determined for each mouse (bottom panel). Each data point represents one mouse and the results were pooled from four independent experiments. **(D)** Quantification of *ex vivo* colonic peristalsis recordings. Peristaltic “clicks” were identified as terminally-oriented waves of colonic muscle contraction. Each data point represents one mouse and the results are pooled from three independent experiments. **(E)** Colonic motility was measured by determining the expulsion time of a glass bead inserted intrarectally into *C1qa*^fl/fl^ and *C1qa*^ΔMφ^ littermates. Each data point represents one mouse and the results are representative of three independent experiments. sm. int., small intestine. Error bars represent SEM. **p<0.01; ***p<0.001; ns, not significant by two-tailed Student’s t-test. **Figure supplements 1 and 2.** *Ex vivo* video recordings of colonic peristalsis in *C1qa*^fl/fl^ and *C1qa*^ΔMφ^ littermates.

## DISCUSSION

Here, we have identified a role for C1q in regulating gastrointestinal motility. We discovered that nerve-adjacent macrophages are the primary source of C1q in the mouse intestine and that macrophage C1q regulates enteric neuronal gene expression and gastrointestinal transit time. Our findings reveal a previously unappreciated function for C1q in the intestine and help to illuminate the molecular basis for macrophage-mediated control of gut motility.

Our study identifies macrophages as the main source of C1q in the mouse small intestine and colon. Both transient antibody-mediated depletion of macrophages and *in vivo* deletion of the *C1qa* gene from macrophages led to a marked reduction in intestinal *C1q* expression. The *C1qa*^ΔMϕ^ mice also lacked C1q in the circulation, indicating that LysM^+^ macrophages or macrophage-like cells are also the source of circulating C1q in the absence of infection. This enhances findings from prior studies indicating that monocytes, macrophages, and immature dendritic cells are the main sources of C1q in the bloodstream (El- Shamy et al., 2018). Importantly, the *C1qa*^ΔMϕ^ mice retained C1q expression in the brain, allowing us to analyze the effects of C1q deficiency without possible confounding effects on the central nervous system.

C1q has two known physiological functions that are distinct and vary according to tissue context. C1q was originally discovered as having a role in the classical complement pathway, which tags and destroys invading microbes (Noris and Remuzzi, 2013; Schifferli et al., 1986). Circulating C1q binds to invading microorganisms and recruits additional proteins that assemble into the membrane attack complex (MAC) (Kishore and Reid, 2000). C1q-mediated MAC formation has been primarily described in the bloodstream, where the necessary accessory proteins are present at high levels (Davis et al., 1979). However, C1q is also expressed in the absence of infection in tissues such as the brain, where it regulates neuronal development and function (Kouser et al., 2015; van Schaarenburg et al., 2016).

Our findings suggest that C1q does not play a central role in immune defense of the intestine. First, we found that intestinal C1q expression was not induced by gut commensals or pathogens and was not deposited into the gut lumen. Second, C1q deficiency did not markedly alter gut microbiota composition or the course of disease after DSS treatment. There were also no major changes in cytokine expression or numbers and frequencies of intestinal immune cells that would indicate dysregulated interactions with the microbiota. Third, C1q was not required for clearance of *C. rodentium*, a non-disseminating enteric pathogen whose clearance requires antigen-specific IgG and complement component 3 (C3) (Belzer et al., 2011). Although we cannot rule out a role for C1q in immune defense against other intestinal pathogens, these findings suggest that C1q is not essential for intestinal immune defense in mice.

Instead, our results indicate that C1q influences enteric nervous system function and regulates intestinal motility. First, most C1q-expressing macrophages were localized to the intestinal submucosal plexus and were largely absent from the lamina propria. Second, C1q-expressing intestinal macrophages were located close to submucosal plexus neurons and expressed cell surface markers like those expressed by nerve-adjacent C1q-expressing macrophages in the lung (Ural et al., 2020). Third, macrophage-specific deletion of *C1qa* altered enteric neuronal gene expression. Finally, consistent with the altered neuronal gene expression, macrophage-specific *C1qa* deletion altered gastrointestinal motility in both the small and large intestine. Thus, our results suggest that the function of C1q in the intestine is similar to its function in the brain, where it regulates the development and function of neurons (Kouser et al., 2015; van Schaarenburg et al., 2016).

A function for macrophage C1q in intestinal motility adds to the growing understanding of how gut macrophages regulate intestinal peristalsis. Prior work has shown that CSF1R^+^ macrophages selectively localize to the muscularis of the mouse intestine (Muller et al., 2014; Gabanyi et al., 2016). These macrophages secrete BMP2, which activates enteric neurons that regulate colonic muscle contraction and thus colonic motility (Muller et al., 2014). We found that depletion of CSF1R^+^ macrophages reduces intestinal C1q expression, and that macrophage-specific deletion of *C1qa* alters enteric neuronal gene expression and gut motility. Thus, our findings suggest that C1q is a key component of the macrophage- enteric nervous system axis.

An important remaining question concerns the molecular mechanism by which C1q regulates gut motility. One possibility is that C1q shapes microbiota composition which, in turn, impacts gut motility. This idea is suggested by studies in zebrafish showing that a deficiency in intestinal macrophages leads to altered gut microbiota composition relative to wild-type zebrafish (Earley et al., 2018). Other studies in zebrafish and mice have shown that severe defects in enteric nervous system development produce changes in gut microbiota composition that are linked to dysregulated gut motility (Rolig et al., 2017; Johnson et al., 2018). However, we did not observe marked changes in the composition of the gut microbiota in *C1qa*^ΔMϕ^ mice, arguing against a central role for the microbiota in C1q-mediated regulation of gut motility. A second possibility is that the absence of C1q leads to immunological defects that alter gut transit time. This idea is consistent with studies showing that T cell cytokines can influence gastrointestinal motility (Akiho et al., 2011). However, this seems unlikely given the lack of pronounced immunological abnormalities in the intestines of *C1qa*^ΔMϕ^ mice.

A third possibility is that C1q interacts directly with enteric neurons or glial cells. Like macrophage-produced BMP2 (Muller et al., 2014), C1q might bind to specific receptors on neurons to regulate their activity. In support of this idea, mouse enteric neurons express *Adgrb1*, which encodes BAI1 (Obata et al., 2020), a known C1q receptor on human neural stem cells (Benavente et al., 2020). Other studies have suggested that C1q functions as a signaling effector on immune cells and neurons (Benoit & Tenner, 2011; Ling et al., 2018). Macrophages have also been reported to restrain neurogenesis in the enteric nervous system through phagocytosis of apoptotic neurons (Kulkarni et al., 2017). Given that C1q can opsonize dying host cells (Botto et al., 1998; Korb and Ahearn, 1997), C1q could regulate enteric nervous system function by tagging apoptotic neurons for elimination. It is also possible that C1q acts directly on enteric smooth muscle cells that regulate gut motility. Although we cannot rule out this possibility, our transcriptional profile of the colonic myenteric plexus of *C1qa*^ΔMϕ^ mice suggests that most of the transcriptional changes were associated with neurons.

Our findings on intestinal C1q have implications for human intestinal disease. Indeed, single cell RNA-seq analysis shows that macrophages recovered from the human intestinal muscularis selectively express *C1q* when compared to lamina propria macrophages (Domanska et al., 2022). Dysregulated peristalsis is a characteristic of irritable bowel syndrome (Vrees et al., 2002) and is present in a subset of inflammatory bowel disease patients (Bassotti et al., 2014). Our finding that macrophage C1q regulates gut motility could suggest new strategies to prevent or treat these diseases. Additionally, most humans with C1q deficiency develop systemic lupus erythematosus (SLE). Since C1q can target cellular debris for phagocytosis, it is thought that C1q deficiency results in increased exposure of self-antigen to the immune system, thereby reducing immune tolerance and causing autoimmune disease (Macedo and Isaac, 2016). Furthermore, roughly 42.5% of SLE patients report gastrointestinal symptoms that range from acute abdominal pain to chronic intestinal obstruction (Fawzy et al., 2016; Tian and Zhang, 2010). The exact cause of these symptoms is unclear. Given that C1q deficiency is strongly correlated with SLE in humans and alters gut motility in mice, we suggest that C1q could be a therapeutic target for SLE patients that present with chronic constipation or other forms of dysregulated intestinal motility.

## MATERIALS AND METHODS

### Key Resources Table

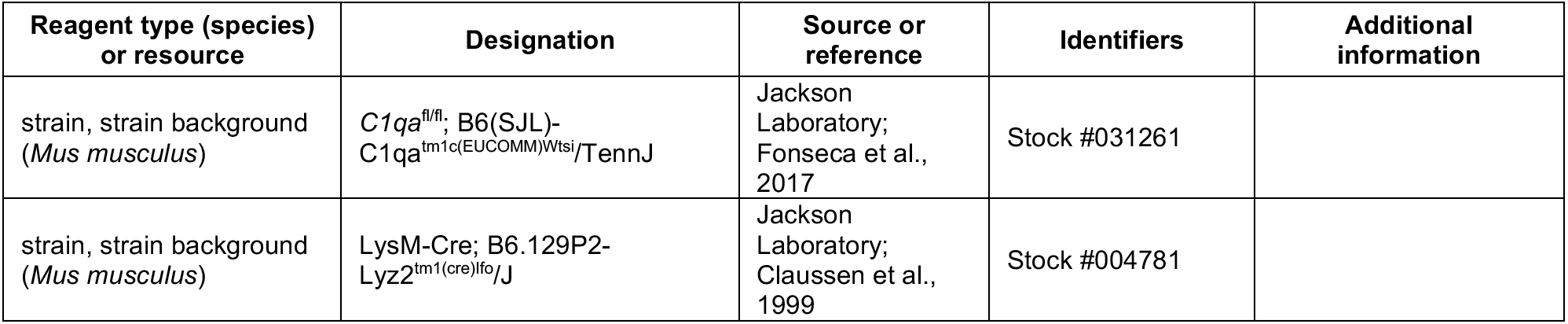
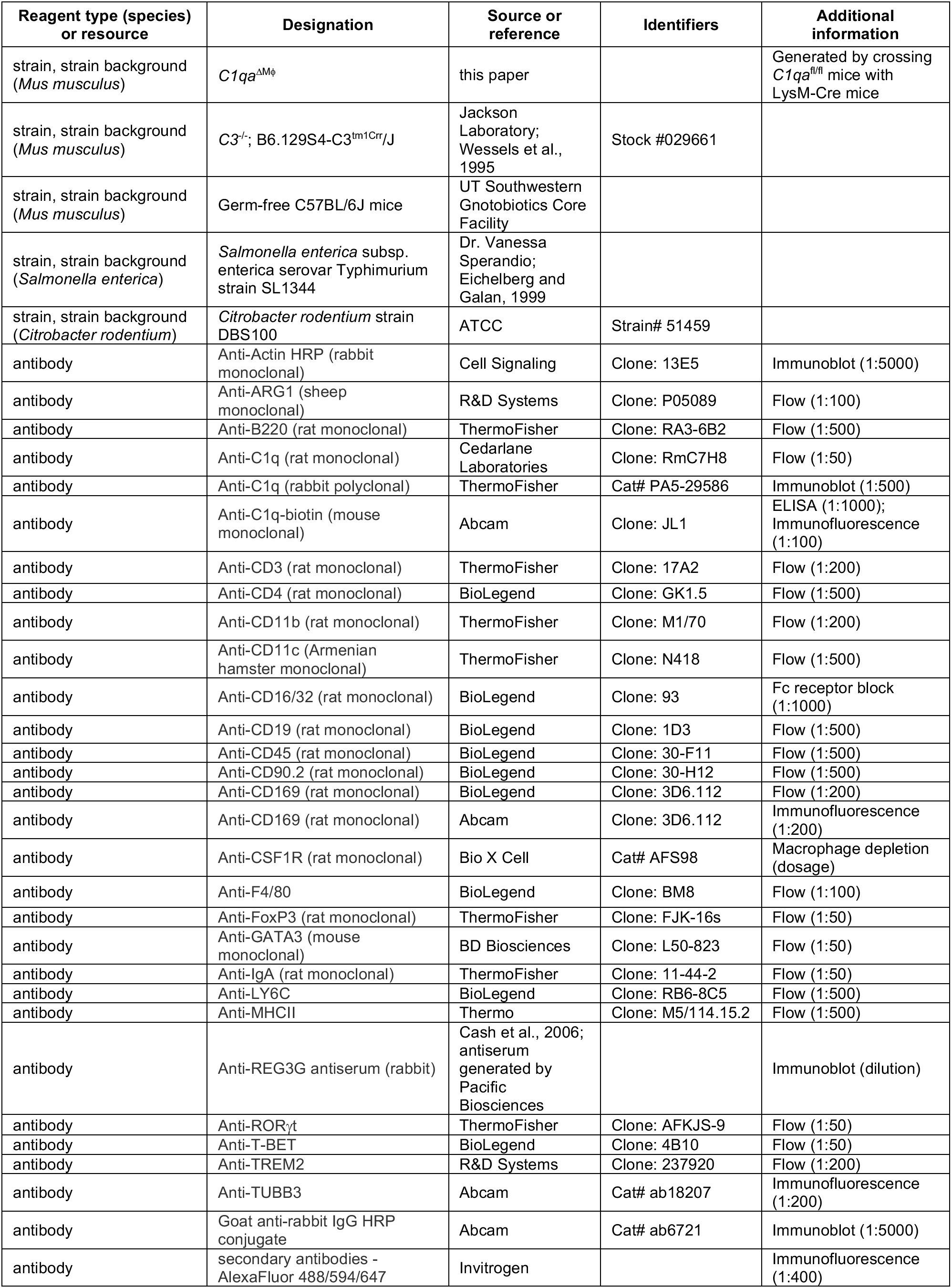
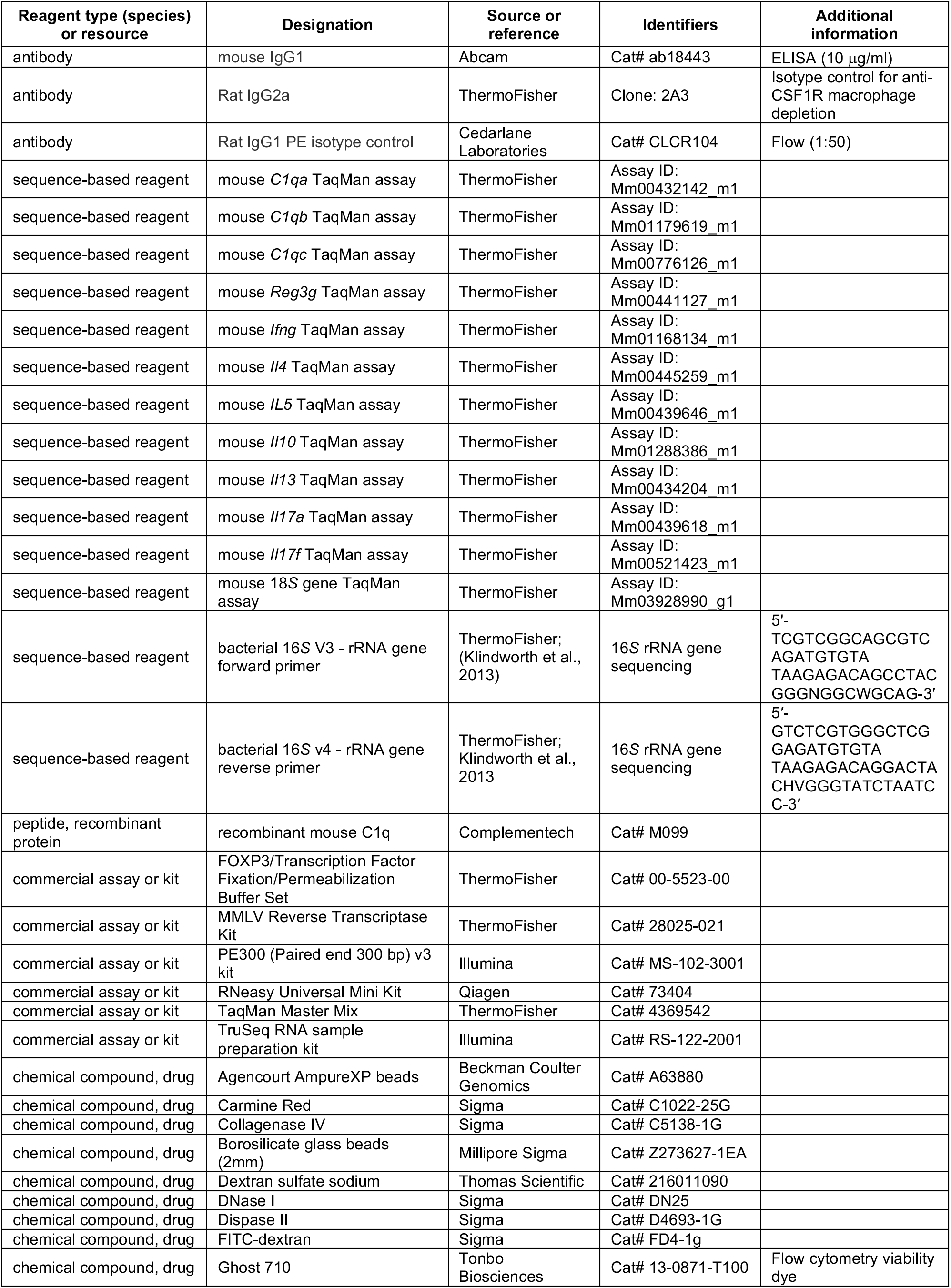
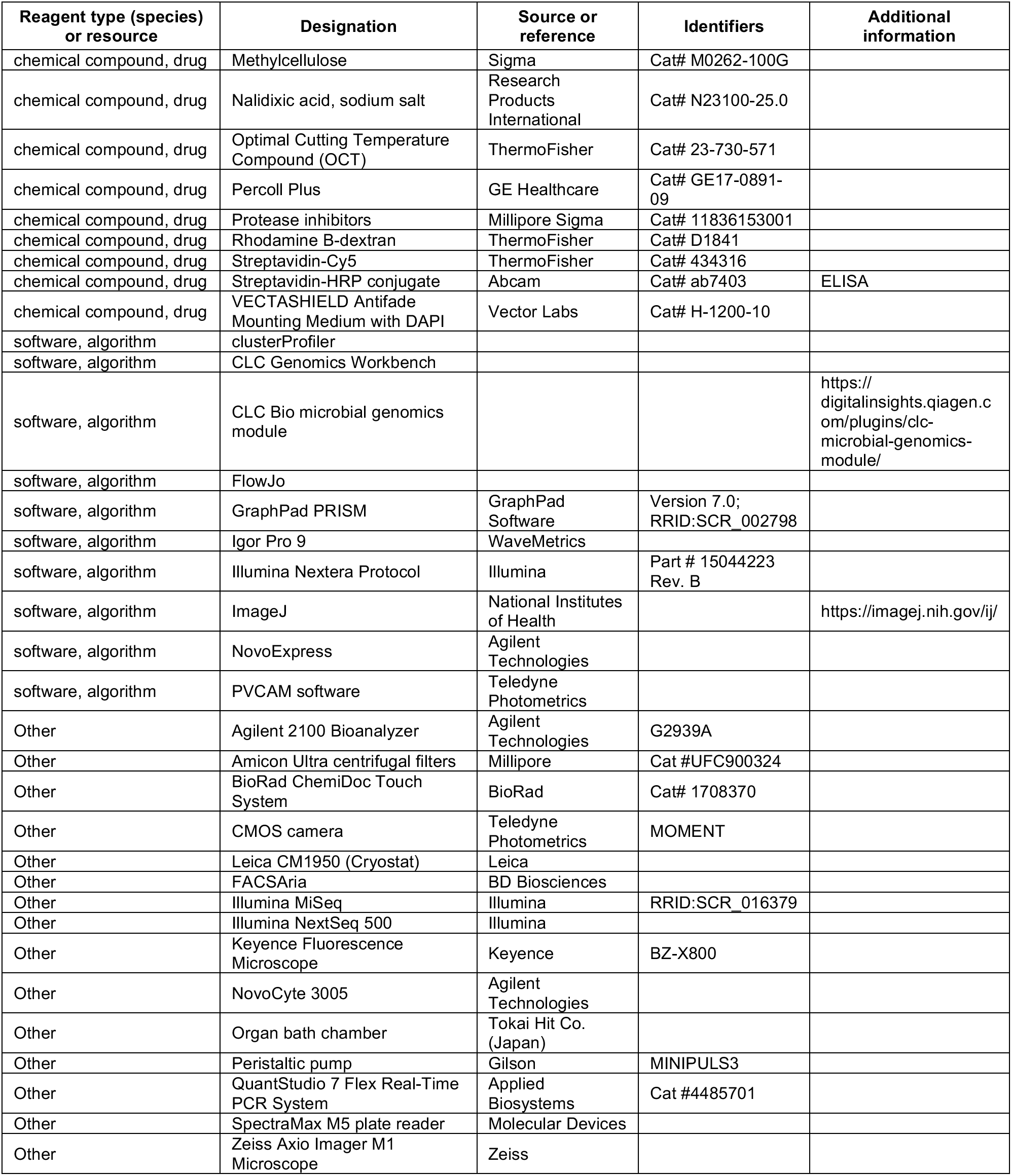

### Mice

Wild-type C57BL/6J (Jackson Laboratory) and *C3^-/-^* mice (Jackson Laboratory; Wessels et al., 1995) were bred and maintained in the SPF barrier facility at the University of Texas Southwestern Medical Center. *C1qa*^ΔMϕ^ mice were generated by crossing *C1qa*^fl/fl^ mice (Jackson Laboratory; Fonseca et al., 2017) with a mouse expressing Cre recombinase controlled by the macrophage-specific mouse *Lyz2* promoter (LysM- Cre mice; Jackson Laboratory; Clausen et al., 1999). Mice 8 to 12 weeks of age were used for all experiments and cohoused littermates were used as controls. Both male and female mice were analyzed. Germ-free C57BL/6J mice were bred and maintained in isolators at the University of Texas Southwestern Medical Center. All procedures were performed in accordance with protocols approved by the Institutional Animal Care and Use Committees (IACUC) of the UT Southwestern Medical Center.

### Quantitative polymerase chain reaction (qPCR)

Tissue RNA was isolated using the RNeasy Universal Mini kit (Qiagen, Hilden, Germany). Cellular RNA was isolated using the RNAqueous Micro kit (ThermoFisher). cDNA was generated from the purified RNA using the M-MLV Reverse Transcriptase kit (ThermoFisher). qPCR analysis was performed using TaqMan primer/probe sets and master mix (ThermoFisher) on a Quant-Studio 7 Flex Real-Time PCR System (Applied Biosystems). Transcript abundances were normalized to 18*S* rRNA abundance. TaqMan probe assay IDs are provided in the Key Resources table.

### Isolation and analysis of intestinal immune cells

Lamina propria (LP) cells were isolated from the intestine using a published protocol (Yu et al., 2013; Yu et al., 2014). Briefly, intestines were dissected from mice and Peyer’s patches were removed. Intestines were cut into small pieces and thoroughly washed with ice-cold phosphate-buffered saline (PBS) containing 5% fetal bovine serum (PBS-FBS). Epithelial cells were removed by incubating intestinal tissues in Hank’s buffered salt solution (HBSS) supplemented with 2 mM EDTA, followed by extensive washing with PBS-FBS. Residual tissues were digested twice with Collagenase IV (Sigma), DNase I (Sigma) and Dispase (BD Biosciences) for 45 minutes at 37°C with agitation. Cells were filtered through 70 μm cell strainers (ThermoFisher) and applied onto a 40%:80% Percoll gradient (GE Healthcare). Subepithelial cell populations were recovered at the interface of the 40% and 80% fractions. For small intestinal cell suspensions, the epithelial fraction was kept and combined with enzymatically liberated subepithelial cells. Cells were washed with 2 mM EDTA/3% FBS in PBS and Fc receptors were blocked with anti-CD16/32 (93). Cells were then stained with the viability dye Ghost 710 (Tonbo Biosciences) followed by antibodies against cell surface markers including anti-CD45 (30-F11), anti-CD11b (M1/70), anti-MHCII (M5/114.15.2), anti-F4/80 (BM8), anti-CD3 (17A2), anti-CD4 (GK1.5), anti-CD19 (1D3), anti-B220 (RA3-6B2), anti-CD11c (N418), anti-CD169 (3D6.112), anti-TREM2 (237920) and anti-LY6C (RB6-8C5). Cells were fixed and permeabilized with the eBioscience FOXP3/Transcription Factor Fixation/Permeabilization buffer set (ThermoFisher) and then subjected to intracellular staining with anti- C1Q (RmC7H8), anti-FOXP3 (FJK-16s), anti-GATA3 (L50), anti-T-BET (4B10), anti-RORγ (AFKJS-9), and anti-ARG1 (P05089). Cells were sorted using a FACSAria (BD Biosciences) or analyzed using a NovoCyte 3005 (Agilent Technologies). Data were processed with FlowJo software (BD Biosciences) or NovoExpress (Agilent Technologies).

### Macrophage depletion

Anti-mouse CSF1R (ThermoFisher; AFS98) and rat IgG2a isotype control (ThermoFisher; 2A3) antibodies were administered intraperitoneally at a concentration of 100 mg/kg. Mice were sacrificed 72 hours post-injection and terminal ileum and colon were collected for qPCR analysis.

### Protein extraction from intestinal cells and feces

To isolate proteins from intestinal cell suspensions, cell pellets were resuspended in 100 μl of RIPA Lysis Buffer (ThermoFisher) supplemented with protease inhibitors (Millipore Sigma) and vortexed vigorously every 5 minutes for 20 minutes. Lysates were cleared of cellular debris by centrifugation at 13000*g* for 5 minutes. To isolate proteins from the intestinal lumen, the entire gastrointestinal tract (from duodenum to distal colon) was recovered from five wild-type C57BL/6J mice. The intestines were flushed with a total of ∼ 50 ml cold PBS containing protease inhibitors (Millipore Sigma, 11836153001). The flushes and fecal pellets were homogenized by rotor and stator (TH Tissue Homogenizer; OMNI; TH01) and large particles were centrifuged at 100*g* for 10 min at room temperature. The supernatants were carefully decanted and centrifuged further at 3000*g* for 20 min at room temperature. The clarified supernatants were precipitated with 40% ammonium sulfate overnight at 4°C. Precipitated protein was centrifuged at 3000*g* for 30 min at 4°C, then resuspended in cold 40% ammonium sulfate and centrifuged again. The pellets were resuspended in room temperature PBS and allowed to mix for 10 min. Protein concentrations were determined by Bradford assay (BioRad).

### Immunoblot

50 μg of fecal protein or 25 μg of cellular protein was loaded onto a 4-20% gradient SDS-PAGE and transferred to a PVDF membrane. Membranes were blocked in 5% nonfat dry milk in Tris-buffered saline (TBS) with 0.1% Tween-20 and then incubated overnight with the following primary antibodies: anti-C1Q (PA5-29586, ThermoFisher) and anti-actin (13E5, Cell Signaling). REG3G was detected by incubating membranes with rabbit anti-REG3G antiserum (Cash et al., 2006). After washing, membranes were incubated with goat anti-rabbit IgG HRP and then visualized with a BioRad ChemiDoc Touch system.

### Enzyme-linked immunosorbent assay (ELISA)

Mouse C1q ELISA was performed as previously described (Petry et al., 2001). Briefly, microtiter plates were coated overnight with mouse IgG1 and were then blocked with 5% BSA in PBS. Serum samples were diluted 1:50 and plated for 1 hour at room temperature. After washing with 0.05% Tween-20 in PBS, bound C1q was incubated with biotinylated anti-C1q antibody (JL1, Abcam). Biotinylated anti-C1q was detected with a streptavidin-HRP conjugate (Abcam). Optical density was measured using a wavelength of 492 nm. Plates were analyzed using a SpectraMax M5 microplate reader (Molecular Devices).

### Intestinal permeability assay

Intestinal permeability assays were performed by treating mice with fluorescein isothiocyanate dextran (FITC-dextran) by oral gavage. The non-steroidal anti-inflammatory drug (NSAID) indomethacin was administered to mice as a positive control. For the experimental group, mice were treated with 190 μl 7% dimethyl sulfoxide (DMSO) in PBS by oral gavage. For the positive control group, mice were treated with 190 μl indomethacin (1.5 mg/ml in 7% DMSO in PBS) by oral gavage. After 1 hour, all mice were treated with 190 μl FITC-dextran (80 mg/ml in PBS) by oral gavage. Mice were sacrificed after 4 hours and sera were collected. Serum samples were centrifuged for 20 minutes at 4°C at 800*g* and supernatants were collected. Serum FITC-dextran levels were measured by a fluorescence microplate assay against a standard curve using a Spectramax plate reader (Molecular Devices).

### Dextran sulfate sodium (DSS) treatment

Age and sex-matched mice were provided with 3% dextran sulfate sodium (weight/volume) in autoclaved drinking water for 7 days. Animal weight and health were monitored in accordance with institutional IACUC guidelines.

### *Salmonella* Typhimurium infection

To prepare bacteria for infection, *Salmonella enterica* serovar Typhimurium (SL1344) was cultured in Luria-Bertani (LB) broth containing 50 μg/ml streptomycin in a shaking incubator at 37°C (Eichelberg and Galan, 1999). The overnight culture was diluted the next day and grown to mid-log phase (OD_600_ = 0.3- 0.5). *C1qa*^fl/fl^ and *C1qa*^ΔMϕ^ littermates were inoculated intragastrically with 10^9^ CFU. All mice were sacrificed 24 hours later and small intestinal tissues were harvested for analysis.

### Citrobacter rodentium infection

To prepare bacteria for infection, an overnight culture of *C. rodentium* (DBS100, ATCC) in LB broth containing nalidixic acid (100 μg/ml) in a shaking incubator at 37°C. The overnight culture was diluted the next day and grown to mid-log phase (OD_600_ = 0.4-0.6). Bacteria were pelleted, washed, and resuspended in PBS. Sex-matched littermates were inoculated intragastrically with 5 × 10^8^ CFU. Fecal pellets were collected at a fixed time every 48 hours, homogenized in sterile PBS, diluted, and plated on LB agar with nalidixic acid (100 μg/ml).

### Immunofluorescence analysis of mouse intestines

Mouse small intestines and colons were flushed with PBS and embedded with Optimal Cutting Temperature compound (OCT) (ThermoFisher). Sections were fixed in ice cold acetone, blocked with 1% BSA, 10% FBS, 1% Triton X-100 in PBS, and then incubated overnight at 4°C with the following antibodies: mouse anti-C1q biotin (JL-1), rat anti-CD169 (3D6.112), and rabbit anti-TUBB3 (ab18207, Abcam). Slides were then washed with PBS containing 0.2% Tween-20 (PBS-T) and incubated with donkey anti-rabbit AlexaFluor 488, donkey anti-rat AlexaFluor 594, and Streptavidin-Cy5 (ThermoFisher) for 1 hour at room temperature in the dark. Slides were then washed in PBS-T and mounted with DAPI-Fluoromount-G (Southern Biotech). Mounted slides were cured overnight at 4°C until imaging.

### RNA-seq analysis of colonic longitudinal muscle myenteric plexus

The colonic longitudinal muscle-myenteric plexus was collected from five age and sex matched *C1qa*^fl/fl^ and *C1qa*^ΔMϕ^ mice by manual dissection using a 2 mm metal probe (Fisher Scientific). RNA was isolated using the RNeasy Mini kit according to the manufacturer’s protocol (Qiagen). Quantity and quality of RNA samples was assessed on an Agilent 2100 Bioanalyzer (Agilent Technologies). RNA-seq libraries were prepared using the TruSeq RNA sample preparation kit (Illumina) according to the manufacturer’s protocol. Libraries were validated on an Agilent Bionalyzer 2100. Indexed libraries were sequenced on an Illumina NextSeq550 for single-end 75 bp length reads. CLC Genomics Workbench 7 was used for bioinformatics and statistical analysis of the sequencing data. The approach used by CLC Genomics Workbench is based on a method developed previously (Mortazavi et al., 2008). To identify differentially enriched biological pathways, all genes were ranked based on their log_2_fold-change and pathway enrichment was identified using the R packages “clusterProfiler” and “msigdbr”. For analysis of differentially expressed genes, gene counts were analyzed using DESeq-2 and differentially expressed genes were defined as having an adjusted p value <0.05. A Fisher’s Exact Test was conducted to assess the overlap between differentially expressed genes in *C1qa*^ΔMϕ^ mice and the TashT mouse (Bergeron et al., 2015).

### 16S rRNA gene sequencing and analysis

The hypervariable regions V3 and V4 of the bacterial 16*S* rRNA gene were prepared using the Illumina Nextera protocol (Part # 15044223 Rev. B). An amplicon of 460 bp was amplified using the 16*S* Forward Primer and 16*S* Reverse Primer as described in the manufacturer’s protocol. Primer sequences are given in the Key Resources Table. The PCR product was purified using Agencourt AmpureXP beads (Beckman Coulter Genomics). Illumina adapter and barcode sequences were ligated to the amplicon to attach them to the MiSeqDx flow cell and for multiplexing. Quality and quantity of each sequencing library were assessed using Bioanalyzer and picogreen measurements, respectively. Libraries were loaded onto a MiSeqDX flow cell and sequenced using the Paired End 300 (PE300) v3 kit. Raw fastq files were demultiplexed based on unique barcodes and assessed for quality. Samples with more than 50,000 quality control pass sequencing reads were used for downstream analysis. Taxonomic classification and operational taxonomic unit analysis were done using the CLC Microbial Genomics Module. Individual sample reads were annotated with the Greengene database and taxonomic features were assessed.

### Gastrointestinal motility assays

Motility assays were adapted from previous studies (Luo et al., 2018; Maurer, 2016; Muller et al., 2014). To determine transit time through the entire gastrointestinal tract, mice were fasted overnight and water was removed 1 hour prior to the start of the experiment. Mice were then singly housed for 1 hour and then gavaged with 100 μl of carmine red (5% weight/volume; Sigma) in 1.5% methylcellulose. Fecal pellets were collected every 15 minutes and transit time was recorded when the dye was first observed in the feces.

For small intestinal motility measurements, mice were fasted overnight and then gavaged with 100 μl of rhodamine B-dextran (5 mg/ml; ThermoFisher) in 2% methylcellulose. After 90 minutes, mice were sacrificed and their stomachs, small intestines, ceca, and colons were collected. Small intestines were cut into 8 segments of equal length and colons were cut into 5 segments of equal length. Segments were cut open lengthwise and vortexed in 1 ml PBS to release rhodamine B-dextran. Fluorescence was then measured on a SpectraMax M5 microplate reader (Molecular Devices). The geometric center of the dye was calculated as: GC = Σ (% of total fluorescent signal per segment × segment number). Relative fluorescence per segment was calculated as: (fluorescence signal in segment / total fluorescence recovered) x 100.

To measure colonic motility, mice were fasted overnight and lightly anesthetized with isoflurane. A 2 mm glass bead was inserted 2 cm intrarectally using a 2 mm surgical probe. Mice were then returned to empty cages and time to reappearance of the bead was recorded.

To account for potential circadian differences in gut motility, the time of day for the initiation of all experiments was held constant.

### *Ex vivo* peristaltic imaging

*Ex vivo* video imaging and analysis of colonic peristalsis was carried out as described previously (Obata et al., 2020). Colons were dissected, flushed with sterile PBS, and pinned into an organ bath chamber (Tokai Hit, Japan) filled with high glucose Dulbecco’s Modified Eagle Medium (DMEM). DMEM was oxygenated (95% O_2_ and 5% CO_2_), run through the chamber using a peristaltic pump (MINIPULS 3, Gilson) and kept at 37°C. Colons were allowed to equilibrate to the organ chamber for 30 minutes before video recording. Movies of colonic peristalsis were captured with a camera (MOMENT, Teledyne photometrics) using PVCAM software (500ms time-lapse delay) over 15 minutes. Movies consist of 3000 sequential image frames that were stitched together in ImageJ and analyzed at a rate of 7 frames per second.

### Statistical analysis

Graphed data are presented as means ± standard error of the mean (SEM). Statistics were determined with GraphPad Prism software. Statistical analyses were performed using two-tailed Student’s t test when comparing two groups, one way ANOVA when comparing multiple groups, and Fisher’s exact test to assess overlap between groups of differentially expressed genes. The statistical tests used are indicated in the figure legends. \**P* ≤ 0.05; \*\**P* ≤ 0.01; \*\*\**P* ≤ 0.001; and ns, *P* > 0.05.

## Acknowledgements

We thank Shai Bel for assistance with immunofluorescence imaging experiments, the UT Southwestern Genomics Core for assistance with RNA sequencing experiments, and the UT Southwestern Flow Cytometry Core for assistance with flow cytometry experiments. *Citrobacter rodentium* strain DBS100 was a gift from Vanessa Sperandio. This work was supported by NIH grants R01 DK070855 (L.V.H.), Welch Foundation Grant I-1874 (L.V.H.), the Walter M. and Helen D. Bader Center for Research on Arthritis and Autoimmune Diseases (L.V.H.), and the Howard Hughes Medical Institute (L.V.H.). M.P. and A.A.C. were supported by NIH T32 AI005284. E.K. was supported by NIH F31 DK126391. Y.O. is the Nancy Cain Marcus and Jeffrey A. Marcus Scholar in Medical Research, in Honor of Dr. Bill S. Vowell.

## ADDITIONAL INFORMATION

### Author contributions

Mihir Pendse, Conceptualization, Investigation, Data curation, Formal analysis, Methodology, Supervision, Writing – original draft, Writing – review and editing; Yun Li, Investigation; Cristine N. Salinas, Investigation; Gabriella Quinn, Data curation, Formal analysis; Daniel C. Propheter, Investigation, Writing – review and editing; Chaitanya Dende, Investigation, Writing – review and editing; Alexander A. Crofts, Data curation, Formal analysis; Eugene Koo, Methodology; Brian Hassell, Investigation; Kelly A. Ruhn, Investigation; Prithvi Raj, Investigation, Data curation, Formal analysis; Yuuki Obata, Investigation, Methodology, Writing – original draft, Writing – review and editing; Lora V. Hooper, Conceptualization, Funding acquisition, Supervision, Writing – original draft, Writing – review and editing.

### Data availability

16*S* rRNA gene sequencing data (***Figure 3D***) and RNA sequencing data (***Figure 4E and F***; ***Figure 4 – figure supplement 2***) are available from the Sequence Read Archive under BioProject ID PRJNA793870. All mouse strains used are available commercially.

---

**Figure 1 - figure supplement 1.**
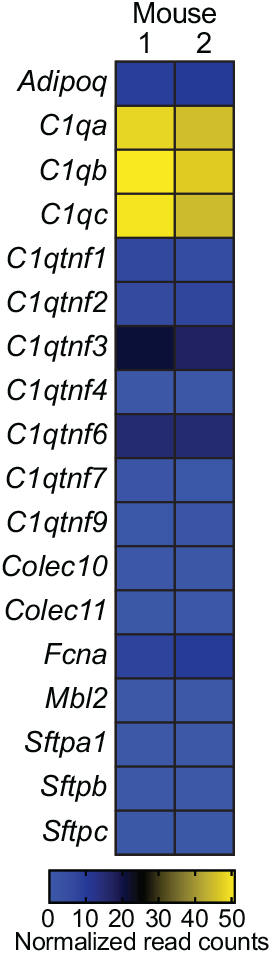
C1q is expressed in the mouse colon. RNA-seq analysis of soluble defense collagen expression in the colons of C57BL/6 mice. Data were reanalyzed from Gattu et al., 2019. Each column represents one mouse. Data are available in the Gene Expression Omnibus repository under accession number GSE122471.

**Figure 2 - figure supplement 1.**
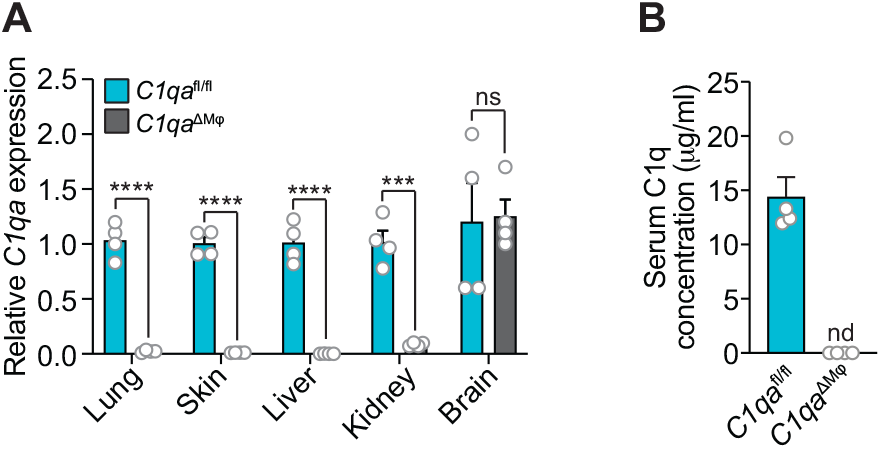
C1q expression is lost systemically but preserved in the central nervous system of *C1qa*^ΔMφ^ mice. **(A)** qPCR measurement of *C1qa* expression in lung, skin, liver, kidney, and brain. Each data point represents one mouse. Data are representative of two independent experiments. **(B)** C1q is absent from the serum of *C1qa*^ΔMφ^ mice. Enzyme-linked immunosorbent assay (ELISA) detection of serum C1q protein from *C1qa*^fl/fl^ and *C1qa*^ΔMφ^ littermates. Data are presented as C1q serum concentration based on a standard curve generated from purified recombinant mouse C1q. Each data point represents one mouse. nd, not detected. Data are representative of three independent experiments. Error bars represent SEM. ***p<0.001; ****p<0.0001; ns, not significant by two-tailed Student’s t-test.

**Figure 3 - figure supplement 1.**
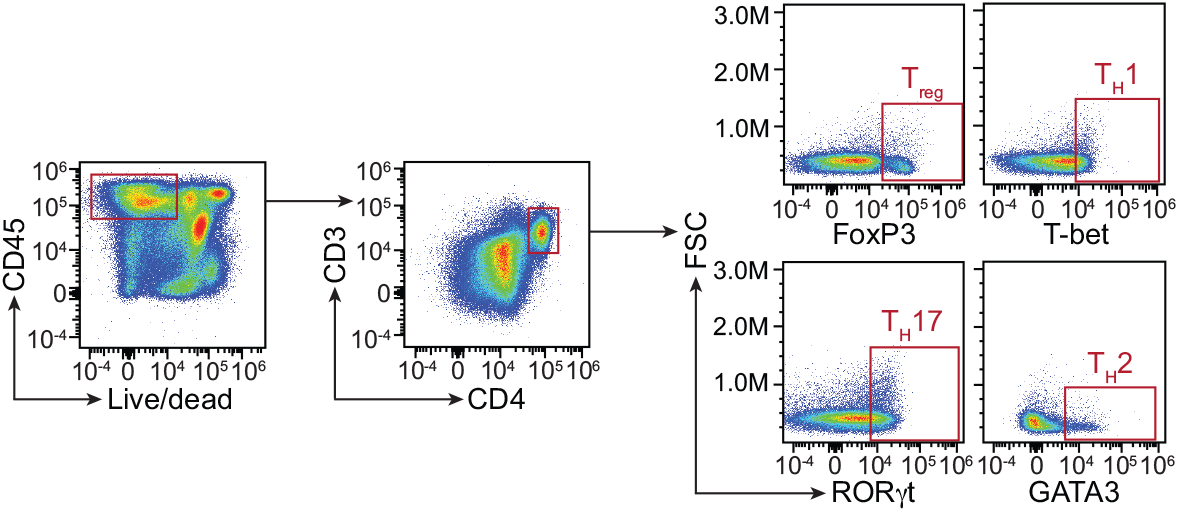
Flow cytometry gating strategy for comparison of T cell populations in *C1qa*^fl/fl^ and *C1qa*^ΔMφ^ mice. Small intestinal cells were recovered from *C1qa*^fl/fl^ and *C1qa*^ΔMφ^ littermates and analyzed by flow cytometry. T cells were gated as live CD45^+^ CD3^+^ CD4^+^. T cell subsets were further identified by gating into T_reg_ (FoxP3^+^), T_H_1 (T-bet^+^), T_H_17 (RORγt^+^), and T_H_2 (GATA3^+^). Representative plots from *C1qa*^fl/fl^ mice are presented and comparisons between *C1qa*^fl/fl^ and *C1qa*^ΔMφ^ littermates are shown in Figure 3. FSC, forward scatter.

**Figure 3 - figure supplement 2.**
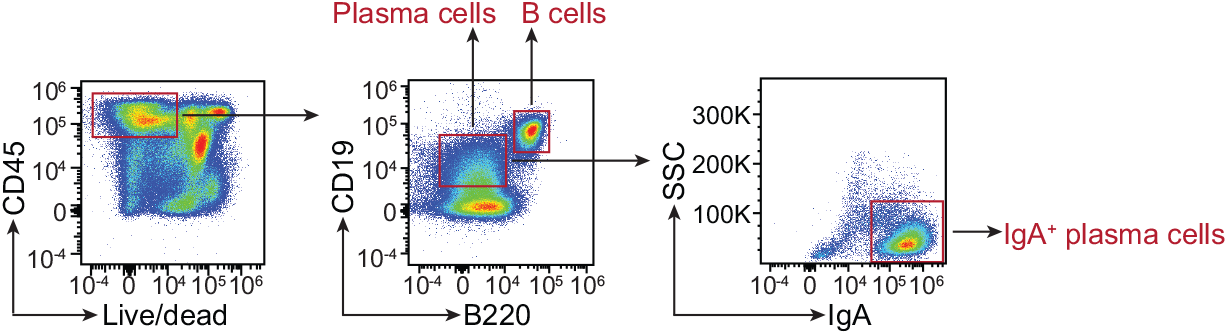
Flow cytometry gating strategy for comparison of B cell and plasma cell populations in *C1qa*^fl/fl^ and *C1qa*^ΔMφ^ mice. Small intestinal cells were recovered from *C1qa*^fl/fl^ and *C1qa*^ΔMφ^ littermates and analyzed by flow cytometry. B cells were gated as live CD45^+^ CD19^+^ B220^+^. Plasma cells were gated as CD19^+^ B220^-^, and IgA^+^ plasma cells were further identified. Representative plots from *C1qa*^fl/fl^ mice are presented and comparisons between *C1qa*^fl/fl^ and *C1qa*^ΔMφ^ littermates are shown in Figure 3. SSC, side scatter.

**Figure 3 - figure supplement 3.**
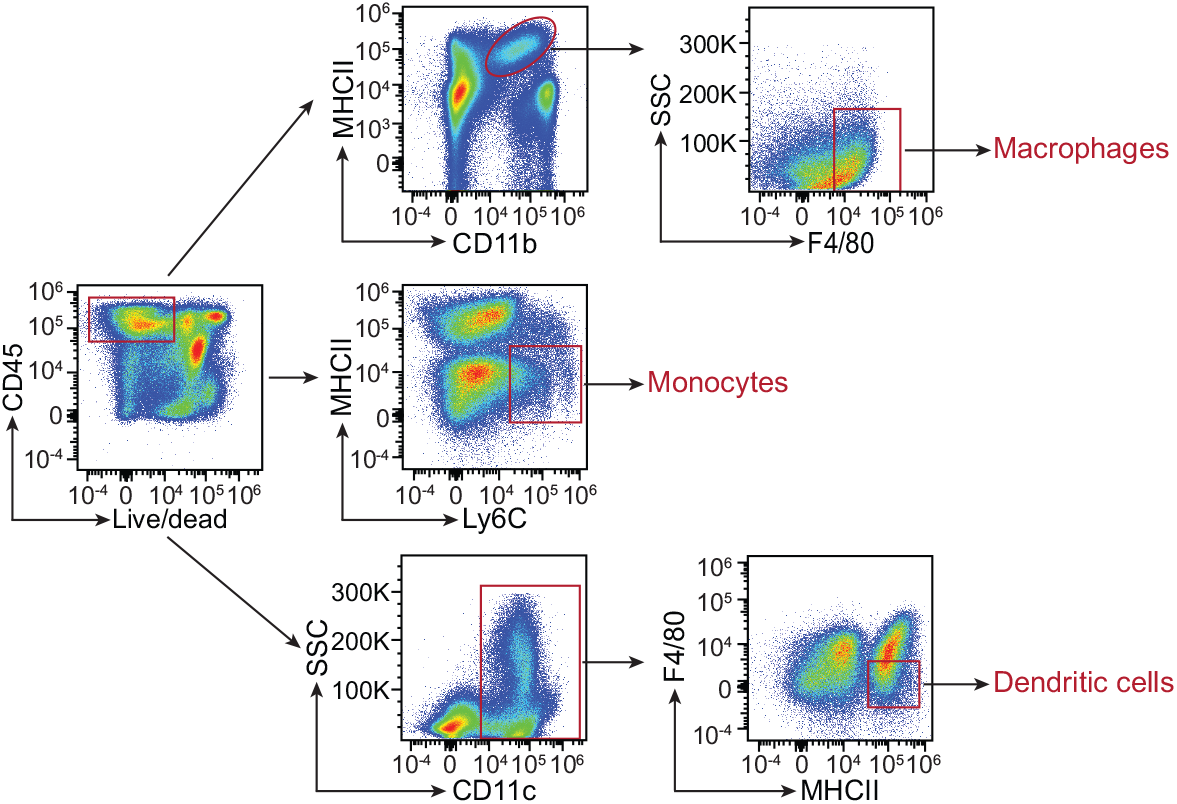
Flow cytometry gating strategy for comparison of myeloid cell populations in *C1qa*^fl/fl^ and *C1qa*^ΔMφ^ mice. Small intestinal cells were recovered from *C1qa*^fl/fl^ and *C1qa*^ΔMφ^ littermates and analyzed by flow cytometry. Macrophages were gated as live CD45^+^ MHCII^+^ CD11b^+^ F4/80^hi^. Monocytes were gated as live CD45^+^ MHCII^-^ Ly6C^+^. Dendritic cells were gated as live CD45^+^ CD11c^+^ MHCII^+^ F4/80^lo^. Representative plots from *C1qa*^fl/fl^ mice are presented and comparisons between *C1qa*^fl/fl^ and *C1qa*^ΔMφ^ littermates are shown in Figure 3. SSC, side scatter; MHCII, major histocompatibility complex II.

**Figure 3 - figure supplement 4.**
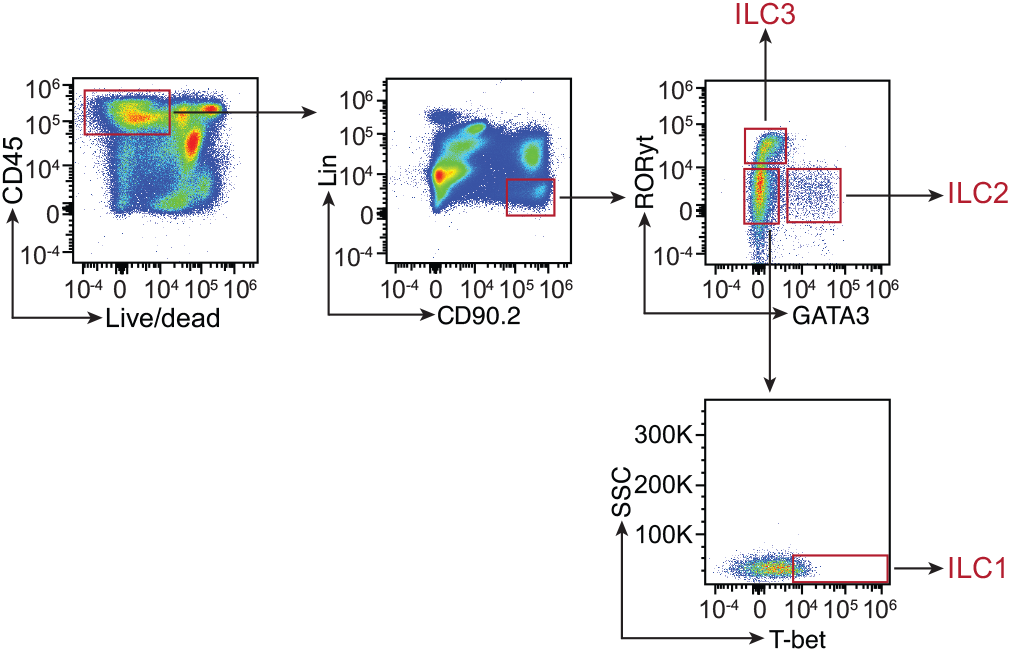
Flow cytometry gating strategy for comparison of innate lymphoid cell populations in *C1qa*^fl/fl^ and *C1qa*^ΔMφ^ mice. Small intestinal cells were recovered from *C1qa*^fl/fl^ and *C1qa*^ΔMφ^ littermates and analyzed by flow cytometry. Innate lymphoid cells (ILC) were gated as live CD45^+^ Lin^-^ CD90.2^+^ and then further identified as ILC1 (RORγt- GATA3^-^ T-bet^+^), ILC2 (RORγt^-^ GATA3^+^) and ILC3 (RORγt^+^ GATA3^-^). Representative plots from *C1qa*^fl/fl^ mice are presented and comparisons between *C1qa*^fl/fl^ and *C1qa*^ΔMφ^ littermates are shown in Figure 3. SSC, side scatter.

**Figure 4 - figure supplement 1.**
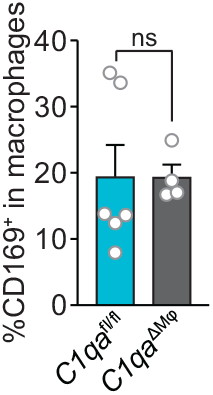
Similar numbers of CD169^+^ macrophages are present in the small intestines of *C1qa*^ΔMφ^ and *C1qa*^fl/fl^ littermates. Flow cytometry analysis of CD169^+^ macrophages was conducted on cells recovered from the small intestines of *C1qa*^ΔMφ^ and *C1qa*^fl/fl^ littermates. CD169^+^ cells were determined as a percentage of total macrophages (live CD45^+^ CD11b^+^ MHCII^+^ F4/80^hi^ cells).

**Figure 5 - figure supplements 1 and 2.**
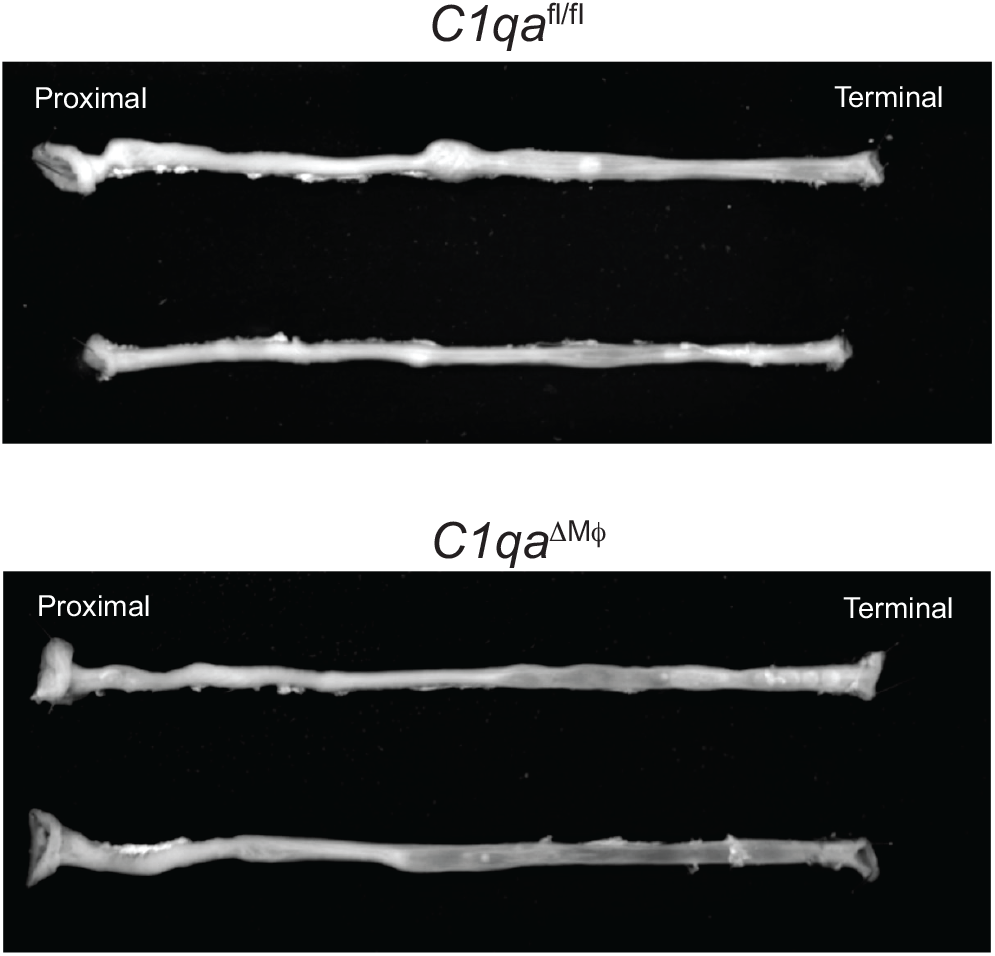
*Ex vivo* video recordings of colonic peristalsis in *C1qa*^fl/fl^ and *C1qa*^ΔMφ^ littermates. Still images captured from *ex vivo* video recordings of colonic peristalsis to indicate orientation of the colons. Colons were allowed to equilibrate to the organ chamber for 30 minutes before video recording. Movies were captured over 15 minutes. Colons from *C1qa*^fl/fl^ mice are shown in figure supplement 1; colons from *C1qa*^ΔMφ^ are shown in figure supplement 2.

